# Opposing effects of an F-box protein and the HSP90 chaperone network on microtubule stability and neurite growth in *Caenorhabditis elegans*

**DOI:** 10.1101/2020.02.13.944967

**Authors:** Chaogu Zheng, Emily Atlas, Ho Ming Terence Lee, Susan Laura Javier Jao, Ken C. Q. Nguyen, David H. Hall, Martin Chalfie

**Author notes:** Correspondence (C.Z.), (M.C.).

## Abstract

Molecular chaperones often work collaboratively with the ubiquitination-proteasome system (UPS) to facilitate the degradation of misfolded proteins, which typically safeguards cellular differentiation and protects cells from stress. In this study, however, we report that the Hsp70/Hsp90 chaperone machinery and an F-box protein, MEC-15, have opposing effects on neuronal differentiation and that the chaperones negatively regulate neuronal morphogenesis and functions. Using the touch receptor neurons (TRNs) of *Caenorhabditis elegans*, we find that *mec-15(−)* mutants display defects in microtubule formation, neurite growth, synaptic development, and neuronal functions, and these defects can be rescued by the loss of Hsp70/Hsp90 chaperones and cochaperones. MEC-15 likely functions in a SCF complex to degrade DLK-1, which is an Hsp90 client protein stabilized by the chaperones. The abundance of DLK-1, and likely other Hsp90 substrates, is fine-tuned by the antagonism between MEC-15 and chaperones; this antagonism regulates TRN development as well as synaptic functions of GABAergic motor neurons. Therefore, a balance between UPS and chaperones tightly controls neuronal differentiation.

**Summary statement:** Molecular chaperones are known to protect cells from stress. However, in this study the authors showed that the Hsp90 chaperone negatively regulates neuronal differentiation when the ubiquitination-proteasome system is compromised.

## Introduction

Molecular chaperones, including the heat shock proteins (Hsps), play essential roles in protein maturation, refolding, and degradation (Hartl et al., 2011; Lindquist and Craig, 1988). Although the function of Hsps in the response to stress has been extensively characterized, their roles in neuronal differentiation are much less understood. Ishimoto et al. (1998) found that Hsp90 promotes neurite extension for the chick telencephalic neurons and spinal neurons *in vitro*. More recently, pharmacological inhibition of Hsp90 by 17-demethoxygeldanamycin (17-AAG) disturbed neuronal polarization and axonal elongation of cultured hippocampal neurons (Benitez et al., 2014). Hsp90 inhibition decreased expression of two Hsp90 client proteins, Akt and GSK3, which have diverse functions in cell differentiation. Thus, Hsp90 may regulate axon specification and growth by affecting specific signaling pathways through its chaperone activity. Given that the Hsp70 and Hsp90 chaperones interact with numerous client proteins, including transcription factors, kinases, and signaling molecules (Wayne et al., 2011), the regulation of neuronal morphogenesis by the chaperones is likely to be context-dependent. Whether Hsp70 and Hsp90 chaperones and their co-chaperones can also negatively regulate neurite growth, however, is unclear.

The ubiquitination-proteasome system (UPS) often works in concert with the chaperone-mediated refolding machinery for protein quality control, which promotes degradation of numerous misfolded proteins in a chaperone-dependent manner (Buchberger et al., 2010). In developing neurons, this process safeguards the protein quality of important guidance molecules. For example, in *Caenorhabditis elegans* the BC-box protein EBAX-1, the substrate-recognition subunit of the Elongin BC-containing Cullin-RING ubiquitin ligase (CRL), and HSP-90/Hsp90 collaboratively regulate the folding and degradation of misfolded SAX-3/Robo receptor during axonal pathfinding (Wang et al., 2013). By preferentially binding to misfolded SAX-3, EBAX-1 not only recruits Hsp90 to promote the refolding of nonnative SAX-3 but also mediates the degradation of irreparable SAX-3 through CRL activity.

Despite known examples of collaboration, UPS and molecular chaperones could theoretically have opposing effects as well, since UPS increases normal protein turnover and chaperones can enhance protein stability. The antagonism of the UPS and the molecular chaperone machinery during neuronal differentiation, however, has not, to our knowledge, been previously reported.

In this study, we find that the F-box and WD40 repeat domain-containing protein MEC-15 (an ortholog of human FBXW9) and ubiquitination promote microtubule (MT) stability and neurite growth by countering the activity of the Hsp70/Hsp90 chaperone network. Mutations in *mec-15* led to a range of developmental defects in the *C. elegans* touch receptor neurons (TRNs), including the loss of microtubules (MTs), inhibited neurite growth and branching, defects in localizing synaptic proteins, and the loss of sensory function. All these defects in *mec-15* mutants can be rescued by removing the Hsp70 and Hsp90 chaperones and co-chaperones, which unexpectedly suggests that the chaperones can disrupt neurite growth and neuronal development and this activity is normally suppressed by the F-box protein and the ubiquitination pathway. Downstream of the chaperones, we identified the MAP3 kinase DLK-1 as an Hsp90 client protein that has MT-destabilizing activity and can inhibit neurite growth. Therefore, our studies provided an important example of how the effects of molecular chaperones are opposed by the ubiquitination pathway during neuronal differentiation.

## Results

### The F-box protein MEC-15 is required for MT stability and neurite growth

The six *C. elegans* touch receptor neurons (ALML/R, PLML/R, AVM, and PVM) are mechanosensory neurons that respond to gentle touch and have a simple morphology (Figure 1A-B). We previously identified a set of neomorphic tubulin mutations that caused the formation of hyperstable MTs and the growth of an ectopic, posteriorly directed neurite in ALM neurons (termed ALM-PN; similarly, ALM-AN is the anteriorly directed neurite of ALM; PLM-AN and PLM-PN are the anteriorly and posteriorly directed neurites of PLM, respectively; Zheng et al., 2017). Among those mutants, the β-tubulin *mec-7(u278;* C303Y) allele had the strongest phenotype, with a very long ALM-PN extending to the tail region. By suppressing this *mec-7(neo)* phenotype, we sought to identify genes that regulate MT stability and neurite development. This *mec-7(u278)* suppressor screen (see Materials and Methods) yielded a *mec-15* nonsense allele *u1042(*R26*), which completely suppressed the growth of ALM-PN in *mec-7(u278)* mutants. Another *mec-15* loss-of-function *(lf)* allele *u75(*Q118*) showed similar suppression (Figure 1A-B and 2A). *mec-15* codes for a protein that contains a N-terminal F-box and four WD-40 repeats (Figure 1C; Bounoutas et al., 2009b) and is orthologous to human FBXW9.

**Figure 1.**
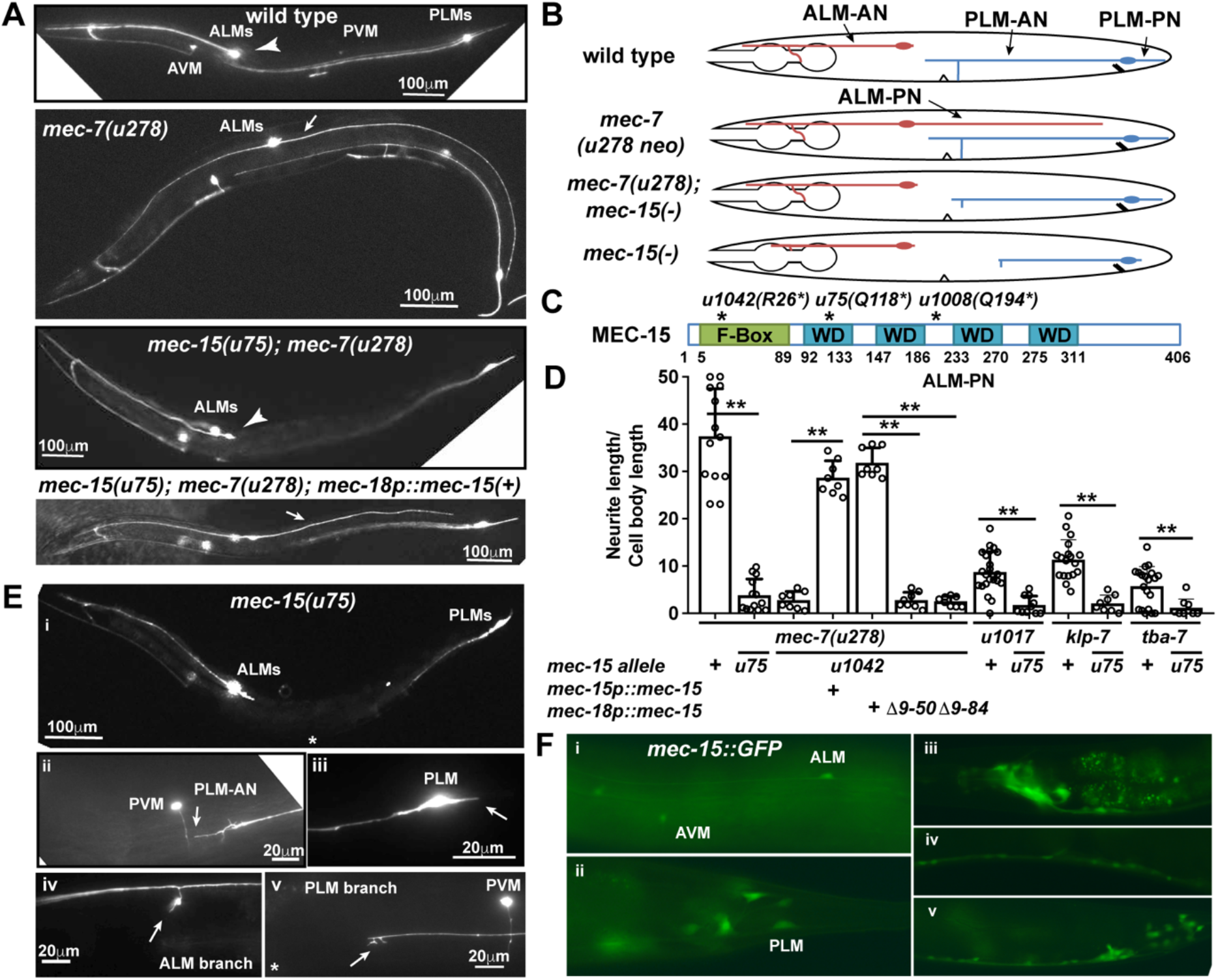
F-box protein MEC-15 promotes neurite growth. (A) The ALM-PN is missing in the wild type animals (arrow head) but is extensive (arrow) in *mec-7(u278 neo)* mutants. *mec-15 lf* mutations suppressed ALM-PN production in *mec-7(u278)* mutants, but the ALM-PN can be restored in the double mutants by expressing wild-type MEC-15 under a TRN-specific *mec-18* promoter. (B) Schematic presentation of ALM and PLM morphologies (with labelled neurites) in wild-type and mutant animals. (C) Protein structures of the wild-type MEC-15 with the F-Box and the four WD-40 repeats (WD) labeled in green and blue, respectively. (D) The normalized length (mean ± SD) of ALM-PN in various strains. Here and in all the other Figures, error bars represent standard deviation, and asterisks indicate significant difference (* for *p* < 0.05 and ** for *p* < 0.01) in ANOVA and Tukey-Kramer test. (E) TRN morphology in *mec-15(u75 lf)* mutants. (i) low magnification view. (ii) PLM-AN (arrow) was shortened and did not extend beyond PVM. (iii) PLM-PN (arrow) was also significantly shortened. Synaptic branches (arrow) of (iv) ALM-AN and (v) PLM-AN could not fully extend in *mec-15* mutants. (F) The expression of *mec-15::GFP* in (i) anterior and (ii) posterior TRNs, (iii) head neurons, (iv) ventral cord neurons, and (v) the preanal ganglion and the tail.

**Figure 2.**
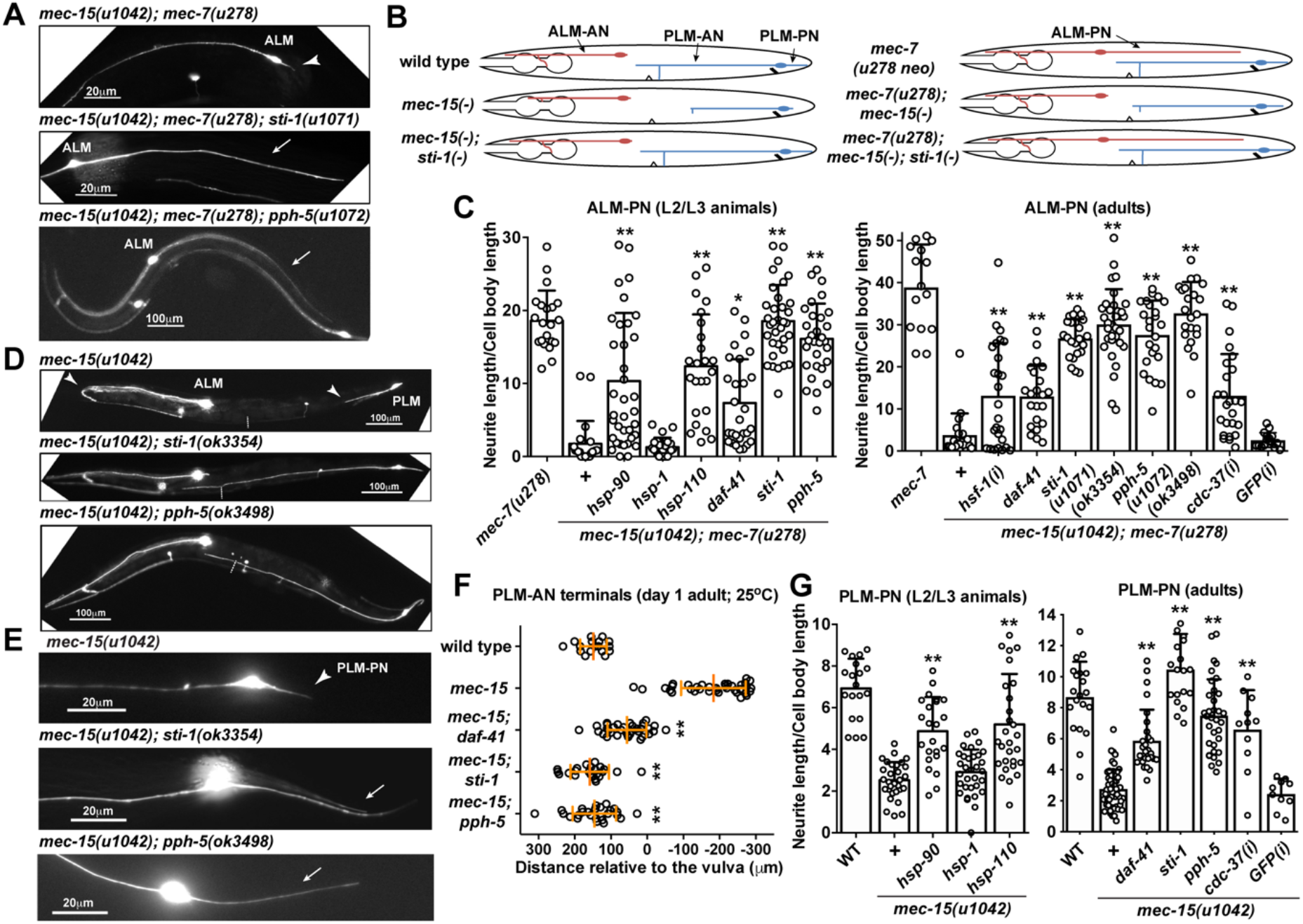
Loss of Hsp90 chaperones suppress the neurite growth defects in *mec-15 lf* mutants. (A) ALM-PN is absent in *mec-15; mec-7* mutants (arrow head) and restored in triple mutants *sti-1* and *pph-5 lf* mutations (arrow). (B) Schematic presentation of ALM and PLM morphologies in wild type and various mutants. *sti-1(−)* is used as an example of *mec-15(−)* suppressors. (C) The length of ALM-PN in various mutants. *hsp-90(ok1333)*, *hsp-1(ok1371)*, and *hsp-110(gk533)* alleles caused larval arrest and were examined at L2/L3 stages. *daf-41(ok3052)*, *sti-1(u1071)*, and *pph-5(u1072)* were examined at both L2/L3 and adult stages; *sti-1(ok3354)* and *pph-5(ok3498)* deletion alleles served as references. *mec-15; mec-7* adult animals carrying transgenes expressing dsRNA against *hsf-1* or *cdc-37* in TRNs were examined; results are labeled as *hsf-1(i)* and *cdc-37(i)*, respectively. (D) Premature termination of ALM-AN and PLM-AN (arrow heads) in *mec-15* single mutants were rescued in *mec-15; sti-1* and *mec-15; pph-5* double mutants. Dashed line indicates the position of the vulva. (E) Shortening of PLM-PN (arrow head) in *mec-15* single mutants were rescued to its normal length (arrows) in the double mutants. (F) Anterior extent of PLM-AN growth in one-day old adults of various mutants grown at 25°C. Distances are given relative to the position of the vulva, so positive values indicate that the PLM-AN grew past the vulva towards the anterior. (G) PLM-PN length in wild type (WT) and various mutants. For (F) and (G), *daf-41(ok3052)*, *sti-1(u1071)*, *pph-5(u1072)*, *hsp-1(ok1371)*, *hsp-90(ok1333)*, and *hsp-110(gk533)* were examined. Asterisks indicate statistical significance for the difference between the *mec-7; mec-15* and the triple mutants (C) or between *mec-15* and the double mutants (G) in ANOVA and Tukey-Kramer test.

The loss of *mec-15* also suppressed the growth of ectopic ALM-PN induced by other mutations that increased MT stability and caused the formation of an ALM-PN (Figure 1D), including another neomorphic β-tubulin mutation *mec-7(u1017;* L377F), a *lf* mutation in a destabilizing α-tubulin gene *tba-7*, and a *lf* mutation in a MT-depolymerizing kinesin-13 gene *klp-7* (Zheng et al., 2017). These data suggest that MEC-15, a F-box protein, is generally required for the excessive neurite growth triggered by elevated MT stability in TRNs.

MEC-15 is also required for normal TRN morphology. *mec-15(lf)* alleles *u75* and *u1042* caused the shortening of both PLM-AN and PLM-PN in an otherwise wild-type background (Figure 1E, 2D, and S1). Another *mec-15* nonsense allele *u1008(*Q194*), isolated from a different screen (Zheng et al., 2017), showed similar TRN morphological defects (Figure S1A-C). Previous studies found that *mec-15* is required for touch sensitivity and the localization of presynaptic proteins in TRNs (Bounoutas et al., 2009b). We found that the ALM-AN and PLM-AN in *mec-15* mutants could not fully extend their synaptic branches (Figure 1E, iv and v), which may have caused the synaptic defects.

As previously found (Bounoutas et al., 2009b; Sun et al., 2013), a *mec-15::GFP* translational reporter was expressed in the TRNs, several head and tail neurons, and ventral cord motor neurons (Figure 1F). We also generated a *mec-15::GFP* knock-in allele at the endogenous *mec-15* locus through CRISPR/Cas9-mediated gene editing, but the resulting strain did not have any detectable GFP expression, suggesting that the endogenous MEC-15 level may be quite low. We next tested whether MEC-15 functions cell-autonomously in the TRNs. Expression of wild-type *mec-15(+)* from the TRN-specific *mec-18* promoter rescued both the neurite growth defects in *mec-15(−)* single mutants and the loss of ALM-PN in *mec-15(−); mec-7(u278 neo)* double mutants. In contrast, the expression of MEC-15 with some or all of the F-box domain removed could not rescue either phenotype (Figure 1D and S1A-C), suggesting that MEC-15 functions within the TRNs and its activity requires the F-box domain.

F-box proteins assemble with Skp and Cullin proteins to form SCF E3 ubiquitin ligase complexes. The involvement of ubiquitination in the action of MEC-15 was supported by the findings that TRN-specific knockdown of *uba-1*, the only known *C. elegans* gene encoding a ubiquitin-activating enzyme, strongly suppressed the growth of ALM-PN in *mec-7(u278 neo)* mutants (Figure S1D). Because mutation or general knockdown of *uba-1* causes sterility and lethality, we performed cell-specific RNAi by expressing double-stranded RNA (dsRNA) against *uba-1* from the TRN-specific *mec-17* promoter. As a negative control, expression of dsRNA against GFP did not affect neurite growth. Moreover, inhibition of proteasomes by bortezomib partially suppressed the growth of ALM-PN (Figure S1D). In addition, previous work found that MT depolymerization led to a general reduction in protein levels in TRNs (Bounoutas et al., 2011). We found that mutations in *mec-15*, knockdown of *uba-1*, and bortezomib treatment all significantly reduced TagRFP expression from a *mec-17p::TagRFP* transgene in the TRNs, measured as fluorescent intensity (Figure S1E). The above data lead to the hypothesis that some MT-destabilizing molecule, which is normally ubiquitinated and targeted for degradation by MEC-15, accumulated abnormally in the *mec-15*-deficient TRNs, leading to reduced MT stability and the inhibition of neurite growth.

Although reduced *uba-1* activity affected ALM-PN outgrowth, we could not identify the other components of the putative SCF complex. In fact, our results suggest that these other components may be redundant. For example, although we found that MEC-15 physically interacted with the Skp homolog SKR-1 in a yeast two-hybrid assay (Figure S1F), *skr-1(ok1696)* null mutations neither caused TRN morphological defects nor suppressed ALM-PN outgrowth in *mec-7(u278 neo)* mutants. In addition, RNAi of *skr-2, skr-8, skr-9, skr-12*, and *skr-13* (other Skp homologs) did not suppress the Mec-7(neo) phenotype (see Materials and Methods for details). Cullin, the third component of the SCF complex, also appeared to be redundant, since mutations or knockdown of Cullin homologs *cul-1*, *cul-2*, *cul-4*, *cul-5*, and *cul-6* did not induce the same defect as the loss of MEC-15.

### Loss of molecular chaperones suppresses neurite growth defects of *mec-15* mutants

To identify the downstream target(s) of the MEC-15-dependent ubiquitination pathway, we screened for suppressors of *mec-15* in the *mec-15(u1042); mec-7(u278)* background and identified the phenotype-causing mutations in eight mutants with restored ectopic ALM-PNs (Table S1; see Materials and Methods for details). Among those mutants were one recessive *lf* allele of *sti-1* and four recessive *lf* alleles of *pph-5*. *sti-1* encodes a homolog of the Sti1/Hop cochaperone that physically links Hsp70 and Hsp90 through the tetratricopeptide repeat (TPR)-domain (Schmid et al., 2012), and *pph-5* encodes a homolog of Protein Phosphatase 5 (PP5), a TPR-domain containing serine/threonine phosphatase that binds to Hsp90 and activates its kinase clients (Vaughan et al., 2008). Deletion alleles of either genes [*sti-1(ok3354)* and *pph-5(ok3498)*] also induced the growth of ALM-PN in *mec-15(u1042); mec-7(u278)* animals (Figure 2A-C and Table S1).

Since STI-1 and PPH-5 are components of the molecular chaperone pathway, we tested other genes in the same pathway. The removal of *hsp-90* (a Hsp90 homolog) and *hsp-110* (a Hsp70 family member) but not *hsp-1* (another Hsp70 family protein) suppressed the loss of MEC-15. *hsp-90(ok1333)* and *hsp-110(gk533)* deletion alleles caused larval arrest, but the arrested *mec-15(u1042); hsp-90(ok1333); mec-7(u278)* and *mec-15(u1042); hsp-110(gk533); mec-7(u278)* triple mutants had long ALM-PNs (Figure 2C).

Moreover, TRN-specific knockdown of *hsf-1/*heat shock transcription factor 1, which activates the expression of Hsps and functions upstream of HSP-90 and HSP-110 (Singh and Aballay, 2006), also induced the growth of ALM-PNs in *mec-15(u1042); mec-7(u278)* double mutants (Figure 2C). PP5, the PPH-5 homolog, dephosphorylates the co-chaperone Cdc37 in yeast and mammals, which is essential for the folding and maturation of Hsp90-dependent protein kinases (Vaughan et al., 2008; Wandinger et al., 2006). TRN-specific RNAi silencing of *cdc-37*, *C. elegans* Cdc37, promoted the growth of ALM-PN in *mec-15(−); mec-7(u278)* animals (Figure 2C), suggesting that some Hsp90 kinase clients may be involved in the regulation of neurite growth.

Through a candidate RNAi screen of 27 Hsp70/Hsp90-related genes (Table S2), we found that knocking down *daf-41*, which encodes the Hsp90 co-chaperone PTGES3/p23, also restored the growth of ALM-PNs in *mec-15(u1042); mec-7(u278)* animals (Figure S2C). A *daf-41* deletion allele, *ok3052*, produced the same phenotype (Figure 2C). Thus, disrupting the Hsp70/Hsp90 chaperone machinery by deleting Hsp70, Hsp90, p23, Sti1/Hop, PP5, or Cdc37 rescued the loss of the F-box protein MEC-15.

Importantly, the suppression of *mec-15* loss did not depend on the presence of *mec-7(u278 neo)*. We crossed *lf* alleles of *hsp-110*, *hsp-90*, *daf-41/p23*, *sti-1*, and *pph-5* with *mec-15(u1042)* to create double mutants and found that the shortening of PLM-PN in *mec-15* single mutants was suppressed in all double mutants (Figure 2E and G). TRN-specific knockdown of *cdc-37* had similar effects (Figure 2G). In addition, we found that the PLM-AN outgrowth defect (but not that of PLM-PN) in *mec-15* null mutants was more severe at 25°C than at 20°C (Figure S2A and S2B), suggesting that the abnormally accumulated proteins may have higher expression or exert a stronger MT-destabilizing activity at higher temperature. This more severe PLM-AN growth defect at 25°C was also suppressed in *mec-15; daf-41*, *mec-15; sti-1*, and *mec-15; pph-5* double mutants; most PLM-ANs extended fully and grew beyond the vulva (Figure 2F). We could not measure the length of PLM-AN in *mec-15; hsp-110* and *mec-15; hsp-90* adults, because those double mutants arrested at early larval stages.

HSP-90 and the cochaperones DAF-41, STI-1, and PPH-5 are expressed in most *C. elegans* cells (Gillan et al., 2009; Richie et al., 2011; Song et al., 2009), and we confirmed their expression in the TRNs (Figure S3A-C). Moreover, we were able to rescue the loss of *daf-41*, *sti-1*, and *pph-5* in their double mutants with *mec-15(u1042)* and their triple mutants with *mec-15(u1042); mec-7(u278)* by expressing the wild-type cochaperone gene from a TRN-specific *mec-17* promoter, indicating that the chaperone pathway genes function cell-autonomously in the TRNs (Figure S3D). Unexpectedly, mutants of chaperones and cochaperones also had strong maternal effects in suppressing the *mec-15*(−) phenotype (Figure S4; see Supplemental Results for details).

Moreover, *hsp-110*, *hsp-90*, *daf-41*, *sti-1*, and *pph-5* single mutants did not show any TRN morphological defects (Figure S5) and *daf-41*, *sti-1*, and *pph-5* single mutants did not cause touch insensitivity (Figure 4G; we could not test *hsp-110* and *hsp-90* homozygous mutants, which are quite uncoordinated and unhealthy and arrest at larval stages). These results suggest that their activities are normally not required for TRN differentiation and function. Nevertheless, an inhibitory role for these Hsp70/Hsp90 chaperones and cochaperones on TRN differentiation is revealed when *mec-15* is lost, because the chaperone proteins were required for the disruption of TRN morphology and functions in *mec-15(−)* mutants.

### Phosphorylation of STI-1 inhibits its activity in regulating neurite growth

Because mutations in *pph-5* and *sti-1* showed the strongest and almost identical effects in suppressing *mec-15(−)* phenotype, we wondered whether, in addition to activating Hsp90 and Cdc37, PPH-5 may also activate STI-1 activity through dephosphorylation. This hypothesis is based on previous findings that the phosphorylation of yeast Sti1 and human Hop, STI-1 homologs, appeared to inhibit its cochaperone activity (Rohl et al., 2015).

Both yeast and human Sti1/Hop proteins contain three TPR domains and two DP (aspartate and proline rich) domains; the TPR2A domain interacts with Hsp90, whereas the TPR1 and TPR2B domains interact with Hsp70 (Schmid et al., 2012). Although *C. elegans* STI-1 contains only TPR2A and TPR2B and one DP domain (Figure 3A), STI-1 is likely capable of transferring clients from Hsp70 and Hsp90. Yeast and human Sti1/Hop proteins contain six and five phosphorylation sites, respectively, which are found mostly flanking the Hsp70-interacting TPR1 and TPR2B domains or in the linker region; those sites are not conserved and appear to be unique to the species (Rohl et al., 2015). Similarly, we predicted potential phosphorylation sites in *C. elegans* STI-1 flanking TPR2B and in the linker region, as well as the only phosphorylatable residue (S289) in the DP domain (Figure 3A); those sites are mostly unique to *C. elegans* STI-1, except Y134 and Y163, which are conserved among yeast, human, and *C. elegans*.

**Figure 3.**
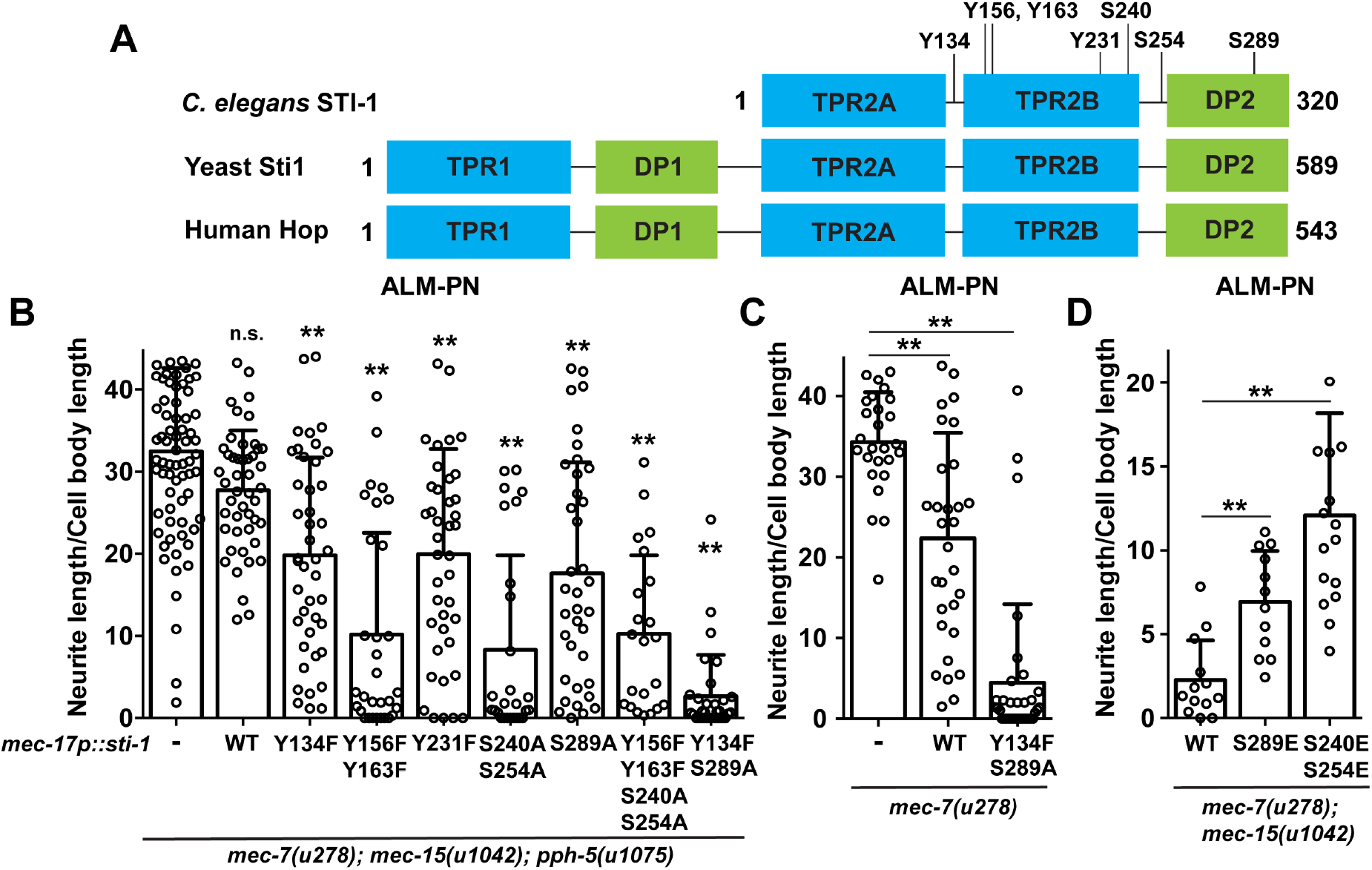
Phosphorylation inhibits STI-1 activity. (A) Schematic representation of domain structure of *C. elegans* STI-1 and its yeast and human homologs. Potential phosphorylation sites for *C. elegans* STI-1 are shown. (B-D) The length of ALM-PN in *mec-7; mec-15; pph-5* triple mutants or *mec-7* single mutants or *mec-7; mec-15* double mutants in the absence of *sti-1* transgene (−) or in the presence of the wild type (WT) or the mutant STI-1 proteins expressed from the *mec-17* promoter. Double asterisks indicate statistical significance (*p* < 0.01) between the control and the test group in ANOVA and Dunnett’s tests.

We reasoned that if PPH-5 regulated STI-1 activity through dephosphorylation, *pph-5(−)* mutants might have hyperphosphorylated STI-1 with low activity and the expression of non-phosphorylatable STI-1 in *pph-5* mutants might rescue STI-1 activity and thus suppress the loss of *pph-5*. Indeed, TRN-specific expression of non-phosphorylatable STI-1 mutants that removed one or several potential modification sites suppressed the effect of *pph-5(−)* mutation to varying degrees in *mec-15(u1042); mec-7(u278)* animals (Figure 3B). As a control, overexpression of wild-type STI-1 had little effect on ALM-PN growth in *pph-5; mec-15; mec-7* animals (Figure 3B). Among the STI-1 mutants, S240A-S254A and Y134F-S289A most strongly suppressed the rescuing effects of *pph-5(−)* mutations on ALM-PN growth, suggesting that 1) those residues are key regulatory sites on STI-1; and 2) PPH-5 functions to activate STI-1. We should point out that PPH-5 is serine/threonine phosphatase and cannot dephosphorylate phospho-tyrosine. So, the inhibitory effect on the tyrosine-to-phenylalanine changes on ALM-PN growth may result from the reduction of overall phosphorylation level of STI-1 and was not specific to the loss of *pph-5*. PPH-5 likely acts through S240, S254, and S289 but not Y134.

Overexpression of STI-1(Y134F-S289A) mutants in *mec-7(u278)* animals also strongly suppressed the growth of ectopic ALM-PN (Figure 3C), indicating that STI-1 dephosphorylation can significantly elevate its activity in destabilizing MTs and inhibiting neurite growth. Consistent with this notion, overexpression of phosphomimic and inactive STI-1 mutants (e.g. S240E-S254E) in *mec-15(u1042); mec-7(u278)* partially suppressed *mec-15(−)* mutation and restored ALM-PN in *mec-15; mec-7* animals presumably by competing with endogenous STI-1 (Figure 3D). The above data indicate that, similar to yeast and mammalian Sti1/Hop proteins, *C. elegans* STI-1 is regulated by inhibitory phosphorylation, and PPH-5 may activate STI-1 through dephosphorylation.

### Mutations in chaperones rescues MT loss and synaptic defects of *mec-15* mutants

Consistent with the hypothesis that chaperones are detrimental for TRN development in *mec-15* mutants, we also observed suppressing effects of chaperone mutations on *mec-15(−)*-induced changes in MT organization. We previously showed that MT organization and stability determined neurite growth patterns in TRNs (Zheng et al., 2017). Using electron microscopy (EM), we found that the number of MTs in a cross section of TRN neurite is dramatically reduced in *mec-15* mutants compared to the wild type in both ALM and PLM (Figure 4A-B). For example, PLM-AN in *mec-15(u1042)* mutants had 4.9 ± 2.9 (mean ± SD) MTs on average in a cross section, compared to 29.8 ± 6.5 MTs in the wild type. In addition to the loss of MTs, *mec-15* mutants also had smaller MTs than the wild type. For example, whereas 98% (N = 306) of the MTs in the wild type had the 15-protofilament (15-p) MTs, 71% (N = 65) of the MTs in the PLM-AN of *mec-15(u1042)* adults had between 11 and 13 protofilaments, resulting in significantly smaller MT diameters (Figure 4C-D). The loss of MTs and the smaller size indicate that MTs are highly unstable and disorganized in the absence of MEC-15.

**Figure 4.**
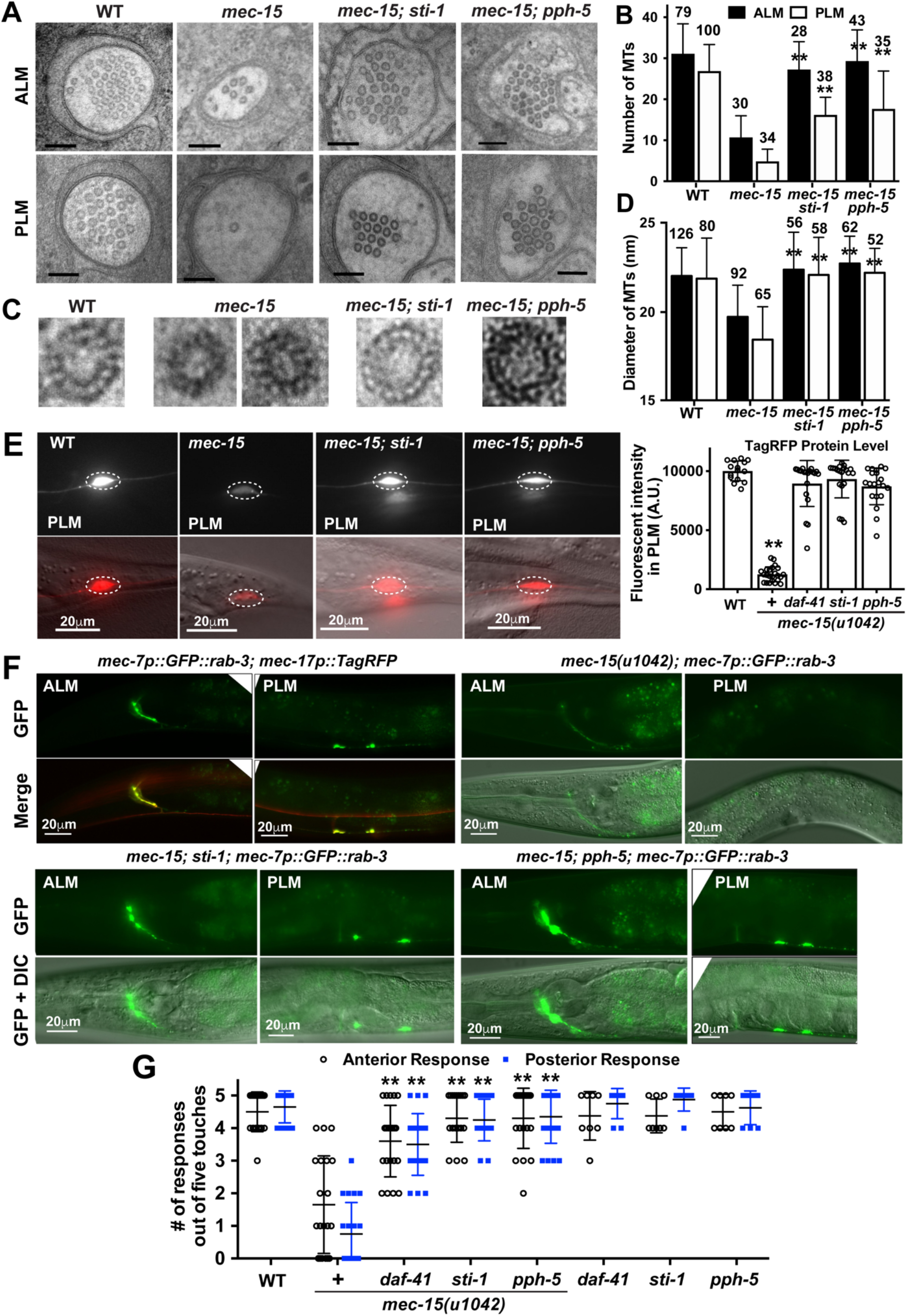
The loss of MTs and TRN developmental and functional defects in *mec-15* mutants were rescued by mutations in Hsp90 cochaperones. (A) Representative EM micrographs of cross-sections of ALM-AN and PLM-AN in wild-type, *mec-15(u1042)*, *mec-15; pph-5(ok3498)*, and *mec-15; sti-1(ok3354)* animals. Scale bar = 100 nm. (B) Average number of MTs in a cross-section of ALM-AN and PLM-AN in various strains. Numbers of sections analyzed for each strain are indicated above the bars. The sections are from at least ten animals. (C) Enlarged EM images of individual MTs with protofilament visualized by tannic acid staining from the cross-sections of PLM-AN of the strains shown in (A). (D) Average diameter of MTs in cross-sections from various strains. Number of MTs measured are indicated above the bars. In (B) and (D), double asterisks indicate statistical significance (*p* < 0.01) for the difference between *mec-15* mutants and the double mutants. (E) Fluorescent intensity of TagRFP in PLM expressed from the *uIs115[mec-17p::TagRFP]* transgene in *mec-15(u1042)*, *mec-15; pph-5(ok3498)*, *mec-15; sti-1(ok3354)*, and *mec-15; daf-41(ok3052)* mutants. Quantification is in arbitrary units for comparison. Representative images are shown with fluorescence only and fluorescence and DIC. (F) Localization of synaptic vesicles labeled by GFP::RAB-3 in TRNs of wild-type, *mec-15*, *mec-15; sti-1*, and *mec-15; pph-5* animals. (G) The number of gentle touch responses from five gentle touch sitmuli in the strains indicated at 20°C. Double asterisks indicate statistical significance (*p* < 0.01) for the difference between *mec-15* mutants and the double mutants.

The structural defects of MTs in *mec-15* mutants are rescued in *mec-15; sti-1* and *mec-15; pph-5* double mutants, which had restored MT numbers, 16.2 ± 4.3 and 17.6 ± 9.2 MTs in the PLM-AN, respectively. The double mutants also have large-diameter 15-protofilament MTs, similar to those seen in the wild type (Figure 4D). These data suggest that the loss of Hsp90 co-chaperones promotes MT formation and stability, which in turn rescues the TRN neurite growth defects in *mec-15* mutants.

Consistent with our previous finding that MT depolymerization in the TRNs causes a general reduction in protein levels (Bounoutas et al., 2011), *mec-15* mutants showed drastically decreased TagRFP expression from a *mec-17p::TagRFP* transgene in the TRNs, compared to the wild type (Figure 4E). This reduction in protein level was completely rescued by mutations in *daf-41/p23*, *sti-1*, and *pph-5* (Figure 4E). Moreover, *mec-15* mutants also showed almost complete loss of GFP::RAB-3 localization at the presynaptic site in PLM neurons and partial loss in ALM neurons (Bounoutas et al., 2009b). This transport defect is likely caused by unstable MTs and is suppressed by the mutations in *sti-1* and *pph-5* (Figure 4F).

TRNs are mechanosensory neurons that detect gentle touch to the body; this sensory function of TRNs depends on the 15-p MTs (Bounoutas et al., 2009a). Mutations in *mec-15* cause touch insensitivity (Chalfie and Au, 1989), likely due to the loss of 15-p MTs. This sensory defect was fully rescued in *sti-1; mec-15* and *pph-5; mec-15* double mutants and partially rescued in *daf-41; mec-15* double mutants (Figure 4G).

Thus, in addition to neurite development, the removal of the molecular chaperone pathway genes restored a variety of MT-related structures and functions affected by *mec-15* loss. Given the protective nature of the chaperones, this restoration is rather unexpected and suggests that Hsp70/Hsp90 machinery may contribute to the refolding and stabilization of a client protein or proteins that negatively affects MT stability and TRN development.

### Hsp90 chaperones inhibit neurite growth by stabilizing DLK-1

Since our *mec-15* suppressor screen (Table S1) yielded two *dlk-1 lf* alleles, *u1105* (V844I) and *u1138* (W394*), we examined the possibility that Hsp70/Hsp90 might regulate MT organization and neurite growth by acting on *dlk-1*, which encodes a MAP3 kinase homologous to human MAP3K12 and MAP3K13. Both *u1105* and *u1138* mutations restored the growth of both ALM-PN and PLM-PN in *mec-15(u1042); mec-7(u278)* mutants (Figure 5A and B). The missense mutation (V844I) in *u1105* is adjacent to a C-terminal region (aa 850-881) of DLK-1 that interacts with its kinase domain (Yan and Jin, 2012); so, catalytic function of DLK-1 may be impaired in *u1105* mutants. Another *dlk-1 lf* allele, *ju476* (a frameshift mutation), suppressed *mec-15(−)* even more strongly. The phenotype of *dlk-1(−)* is similar to the effects of *sti-1(−)* and *pph-5(−)* in *mec-15; mec-7* mutants. Independent of the *mec-7(u278)* background, *dlk-1 lf* alleles also suppressed the shortening of PLM-PN and the premature termination of PLM-AN in *mec-15(u1042)* mutants (Figure 5C and D). Moreover, *dlk-1 lf* alleles showed maternal effects similar to *daf-41*, *sti-1*, and *pph-5 lf* alleles (Figure 5B). These data suggest that *dlk-1* may function in the same pathway as the chaperone and co-chaperone genes.

**Figure 5.**
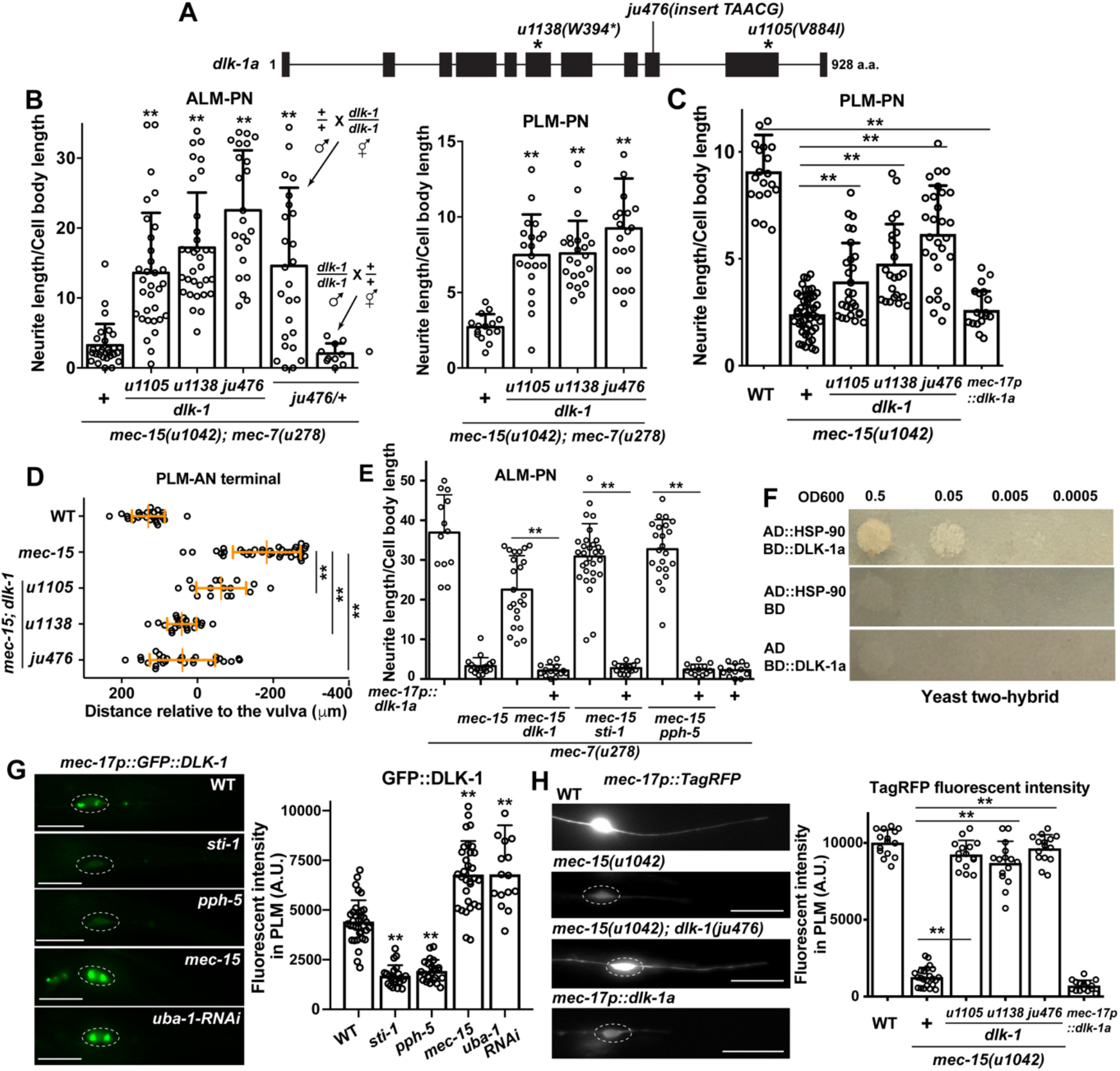
DLK-1 acts downstream of the chaperones to destabilize MTs and inhibit neurite growth. (A) Gene structure of the long isoform (a) of *dlk-1* and the positions of three *lf* mutations. (B) Effects of *dlk-1* mutations on the normalized neurite length of ALM-PN and PLM-PN in *mec-15(u1042); mec-7(u278)* animals. The ALM-PN length in *ju476/+* animals was examined in male cross progeny. Double asterisks indicate significant difference (*p* < 0.01) between the test group and *mec-15; mec-7* double mutants. (C) The length of PLM-PN in *dlk-1; mec-15* mutants and wild-type animals expressing *mec-17p::dlk-1a* transgene. (D) The distance between the PLM-AN terminal and the vulva in the *dlk-1; mec-15* mutants grown at 25°C. (E) The length of ALM-PN in *mec-7(u278)* animals with indicated mutant genotype. Some animals expressed a *mec-17p::dlk-1a* transgene. The *dlk-1(ju476)*, *sti-1(ok3354)*, and *pph-5(ok3498)* alleles were used. (F) Yeast two-hybrid assays for the interaction between AD::HSP-90 and BD::DLK-1a. AD means the Gal4 activation domain and BD means the Gal4 DNA-binding domain. (G) Fluorescent intensity of GFP::DLK-1 expressed from the TRN-specific *mec-17* promoter in wild type, *sti-1(ok3354)*, *pph-5(ok3498)*, and *mec-15(u1042)* mutants and animals carrying *mec-17p::uba-1RNAi* transgenes. Dashed circles enclose PLM cell bodies; scale bar = 20 μm. Double asterisks indicate significant difference from WT. (H) Fluorescent intensity of TagRFP expressed from *mec-17* promoter in *dlk-1; mec-15* double mutants and in wild type animals expressing *mec-17p::dlk-1a* transgene.

Expression of DLK-1a from the TRN-specific *mec-17* promoter rescued the loss of *dlk-1* by suppressing the growth of ALM-PN in *dlk-1; mec-15; mec-7* triple mutants, confirming that DLK-1 functions cell-autonomously (Figure 5E). More importantly, overexpression of DLK-1a also suppressed the phenotype of *pph-5* and *sti-1* mutants, indicating that DLK-1 functions downstream of the chaperones (Figure 5E). Through a yeast two-hybrid assay, we detected a physical interaction between HSP90 and DLK-1a (Figure 5F), which is the long and active isoform of DLK-1 (Yan and Jin, 2012). Moreover, the level of GFP::DLK-1a fusion protein is drastically reduced in *pph-5* and *sti-1* mutants, suggesting that the stability of DLK-1 relies on the Hsp90 chaperones (Figure 5G).

DLK-1 is known to induce MT dynamics and is essential for axonal regeneration in TRNs (Tang and Chisholm, 2016; Yan et al., 2009); DLK-1 signaling is also required for MT depolymerization-induced downregulation of protein levels in TRNs (Bounoutas et al., 2011). Indeed, *dlk-1 lf* mutations blocked the downregulation of TagRFP in *mec-15(−)* animals, while overexpression of DLK-1a in the wild-type animals led to marked reduction of TagRFP expression (Figure 5H). Consistent with a role in reducing MT stability, DLK-1a overexpression strongly suppressed the growth of ectopic ALM-PN in *mec-7(u278)* animals and also shortened PLM-AN and PLM-PN in wild-type animals (Figure 5C and E). The above data suggest that DLK-1 likely mediates the activity of Hsp90 chaperones and cochaperones in destabilizing MTs during development. Coincidentally, a recent study found that Hsp90 is also a chaperone for DLK in mouse neurons and in Drosophila and is required for axon injury signaling (Karney-Grobe et al., 2018). Thus, the regulation of DLK-1 by Hsp90 chaperone appears to be evolutionarily conserved.

Importantly, DLK-1 is not the only downstream effector of the chaperones, because unlike the *sti-1* and *pph-5 lf* mutations, the *dlk-1 lf* mutations did not rescue the touch insensitivity, PLM-AN branching defects, or the GFP::RAB-3 localization defect in *mec-15* mutants (Figure S6). Even for neurite growth, the loss of *dlk-1* did not fully rescue the outgrowth defect of PLM-PN and PLM-AN in *mec-15* mutants (Figure 5C and D). Thus, we expect that other Hsp90 client proteins also function during TRN development.

### MEC-15 downregulates DLK-1 levels in TRNs

Since MEC-15 is a F-box protein that likely functions in a SCF complex to target substrate protein for ubiquitination and degradation, we next asked whether MEC-15 regulates DLK-1 protein levels. We found that the expression of GFP::DLK-1 fusion proteins was elevated in *mec-15* mutants (Figure 5G), whereas the levels of STI-1::GFP and PPH-5::GFP fusion proteins were not changed in *mec-15* mutants (Figure S7A). Importantly, the level of GFP::DLK-1 was also increased by blocking ubiquitination through TRN-specific silencing of *uba-1* (Figure 5G). Thus, MEC-15 likely targets DLK-1 but not the Hsp90 co-chaperones for degradation.

We could not detect any physical interaction of MEC-15 with DLK-1a in yeast two-hybrid assays, suggesting that the interaction may be transient or dependent on particular post-translational modification of DLK-1. We did not detect any interaction of MEC-15 with STI-1, PPH-5, DAF-41, HSP-90, or HSP-110 either. In attempts to detect MEC-15 binding to DLK-1 in the TRNs, we made constructs that express HA-tagged MEC-15 and FLAG-tagged DLK-1 from the TRN-specific *mec-18* and *mec-17* promoters, respectively. When we injected the two expression constructs into worms separately, both MEC-15 and DLK-1 can be detected by western blot probing the tags; but when injected together, only MEC-15::HA was detected, and FLAG::DLK-1 was undetectable (Figure S7B). Although this renders the co-immunoprecipitation assay impossible, these results suggested that MEC-15 suppressed the expression of DLK-1 protein.

RPM-1, a RING-finger E3 ubiquitin ligase, also downregulates the abundance of DLK-1 at the protein level and modulates the p38 MAP Kinase pathway (Nakata et al., 2005). The phenotype of *rpm-1* mutants, however, is the direct opposite of *mec-15* mutants. Instead of the shortened TRN neurites of *mec-15* mutants, *rpm-1(ok364)* knockout mutants had overextended ALM-AN and PLM-AN (Figure S7C; Schaefer et al., 2000). Moreover, the loss of *rpm-1* did not suppress the ectopic ALM-PN in *mec-7(u278)* mutants, and the *rpm-1 mec-7* double mutants showed both long ALM-PN and the overextension of ALM-AN and PLM-AN (Figure S7D-F). Thus, RPM-1 appears to negatively regulate neurite growth, whereas MEC-15 promotes growth. Downstream of both proteins, DLK-1 may exert dual functions in both inducing MT dynamics and promoting neurite extension depending on the cellular contexts.

*mec-15* is epistatic to *rpm-1*, since *mec-15; rpm-1* double mutants showed mostly the *mec-15(−)* phenotype, having shortened PLM-ANs and PLM-PNs. The *mec-15; rpm-1; mec-7* triple mutants showed the phenotype of *mec-15; mec-7* double mutants in the suppression of ALM-PN (Figure S7E-F). This epistasis suggests that either MEC-15-mediated regulation of DLK-1 plays a more dominant role in TRNs than RPM-1-mediated regulation or MEC-15- regulated DLK-1-independent pathways act downstream of RPM-1.

### MEC-15 regulates the function of GABAergic motor neuron by antagonizing Hsp90 chaperone activities

Finally, the genetic interaction between *mec-15* and the Hsp90 chaperones occurs not only in the TRNs, but also in other neurons. In the ventral cord motor neurons, MEC-15 regulates the trafficking of synaptic vesicle protein SNB-1 and promotes GABAergic synaptic transmission (Sun et al., 2013). We found that *mec-15* mutants had fewer synaptic puncta labeled by SNB-1::GFP in GABAergic neurons than wild-type animals (Figure 6A-C). This deficit in synaptic density was rescued by mutations in *sti-1* or *pph-5*, although the average fluorescent intensity of each puncta was similar between the mutants and the wild-type animals (Figure 6A-C).

**Figure 6.**
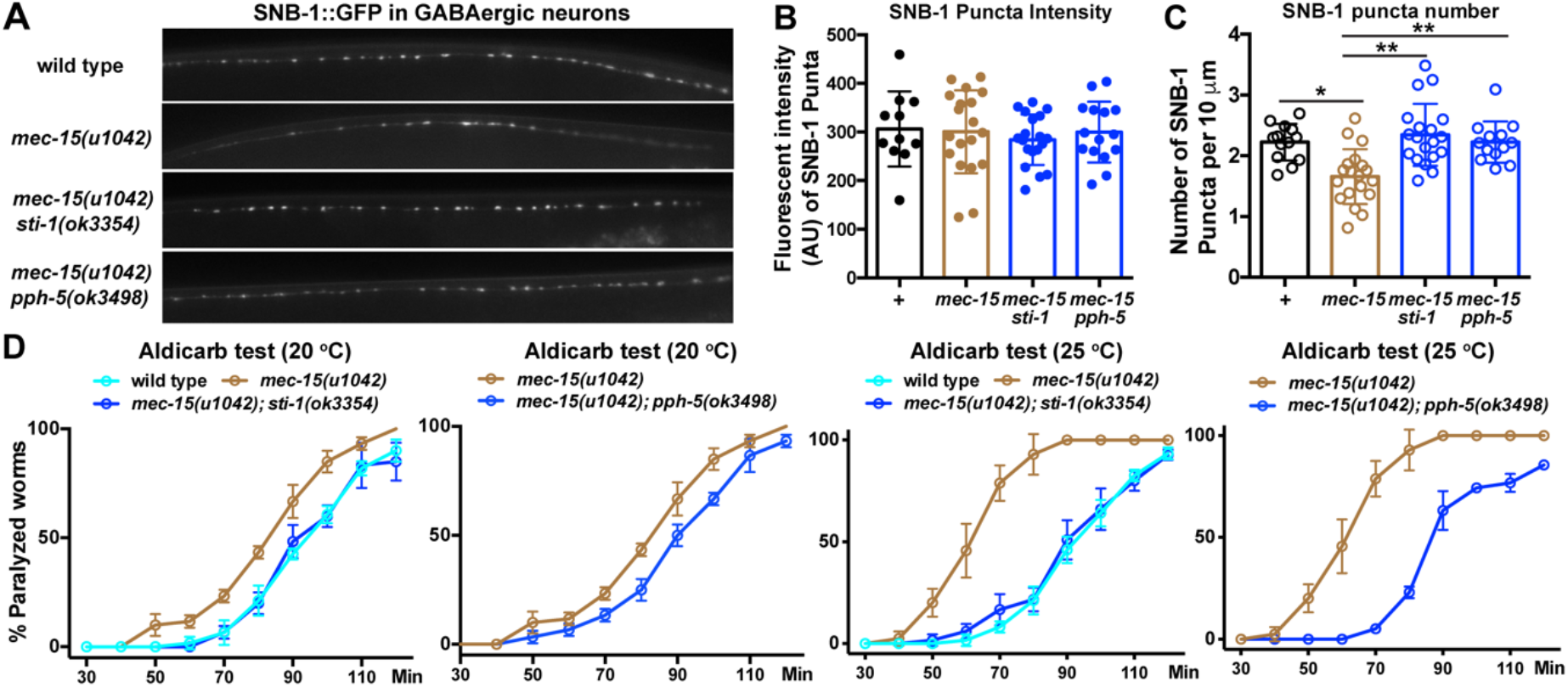
MEC-15 regulates synaptic functions of GABAergic motor neuron by antagonizing the activity of Hsp90 chaperones. (A) Synaptic puncta in the ventral neurite of GABAergic motor neurons labeled by *juIs1[unc-25p::snb-1::GFP]* in the wild-type animal, *mec-15*, *mec-15; sti-1*, and *mec-15; pph-5* mutants. (B) Average fluorescent intensity (arbitrary unit; AU) of the SNB-1::GFP puncta in the indicated strains. (C) Average number of SNB-1::GFP puncta in 10 μm of the ventral neurite of GABAergic motor neurons in the indicated strains. (D) Percentage of adult animals paralyzed at the indicated time point after being exposed to 1 mM Aldicarb. Animals were raised from embryos to young adults at either 20°C or 25°C before test. 20 animals and three replicates were used in each experiment. The experiment was repeated three times; mean ± SD was shown. The same *mec-15* data were plotted twice in the two graphs for comparing with the double mutants.

Behaviorally, reduced GABA release in *mec-15* mutants led to lower inhibitory postsynaptic currents in the body wall muscles and increased sensitivity to the acetylcholine esterase inhibitor aldicarb, which causes paralysis through the accumulation of acetylcholine and persistent muscle contraction (Sun et al., 2013). This increased sensitivity to aldicarb in *mec-15* mutants was completely suppressed in *mec-15; sti-1* and *mec-15; pph-5* mutants (Figure 6D). Therefore, MEC-15 and Hsp90 chaperones also appear to counter each other to regulate GABAergic synaptic function. Moreover, similar to the thermosensitive PLM-AN growth defects in *mec-15* mutants, we found that *mec-15(−)* animals grown at 25°C are much more sensitive to aldicarb than animals raised at 20°C; loss of *sti-1* or *pph-5* also rescued this temperature-dependent defects (Figure 6D). These results suggest that MEC-15 target proteins in both TRNs and GABAergic motor neurons are more stable or have higher levels at higher temperature, likely due to the increased activity of Hsp90 chaperones at higher temperature.

## Discussion

### Ubiquitination and protein degradation promote stable MTs and neurite growth

The ubiquitination-proteasome system (UPS) affects many processes of neuronal development, including neurogenesis, cell fate specification, neuronal migration, polarization, and axonal and dendrite morphogenesis (Tuoc and Stoykova, 2010; Yamada et al., 2013). With respect to neurite morphogenesis, previous studies suggested that E3 ubiquitin ligase can both positively and negatively regulate neurite growth. For example, during axodendritic polarization in hippocampal neurons, the HECT-domain E3 ubiquitin ligase Smurf1 promotes the growth of axons by degrading RhoA (inhibitor of axon growth), whereas Smurf2 inhibits the growth of neurites fated to be dendrites by degrading Ras GTPase Rap1B (inducer of neurite growth; Cheng et al., 2011; Schwamborn et al., 2007).

Our studies identified the F-box and WD40 domain protein MEC-15 (ortholog of human FBXW9) as a positive regulator of neurite growth. As in the above examples, MEC-15 presumably functions in a Skp, Cullin, F-box containing complex (or SCF complex), which is a E3 ubiquitin ligase, to degrade some protein(s) that can inhibit neurite outgrowth. One potential substrate of MEC-15 is the MAP3 kinase DLK-1, which is known to induce MT dynamics (Tang and Chisholm, 2016) and can suppress neurite growth when overexpressed (this study). Since the loss of *dlk-1* did not fully rescue the developmental defects of *mec-15(−)* mutants, MEC-15 likely has multiple substrates.

In the PLM neurons, the anterior neurite is considered an axon because its MTs has uniform “plus-end out” polarity, whereas the posterior neurite is more like a dendrite given that its MTs have mixed polarity (Hsu et al., 2014). MEC-15 is required for the extension of both PLM-AN and PLM-PN, which suggests that MEC-15 is a general regulator of neurite growth instead of an axon- or dendrite-specific regulator, like the E3 ligase Cdh1-APC and Cdc20-APC (Konishi et al., 2004). This function of MEC-15 may be attributed to its activity in promoting the formation of stable MTs, since mutations in *mec-15* led to significant loss of MTs and the reduction of MT diameters. Thus, our studies link SCF complex and UPS to the general stability of MT cytoskeletons.

### Dual yet contradictory functions of DLK-1

Another conserved RING-domain E3 ubiquitin ligase, RPM-1, also regulates neurite extension in TRNs by interacting with the SCF complex, including SKR-1/Skp, CUL-1/Cullin, and the F-box protein FSN-1/FBXO45 (Nakata et al., 2005). However, MEC-15 and RPM-1 have opposing functions. MEC-15 promotes neurite growth, whereas RPM-1 inhibits neurite growth. More surprisingly, both RPM-1 and MEC-15 appear to target DLK-1 for degradation. Since mutations in *dlk-1* partially suppress both the overextension phenotype in *rpm-1(−)* mutants (Nakata et al., 2005) and the underextension in *mec-15(−)* mutants (this study), DLK-1 may exhibit dual yet opposite functions in regulating neurite growth.

Elevated DLK-1 level promotes axon growth in *rpm-1(−)* mutants but inhibits neurite growth in *mec-15(−)* mutants; these two opposing functions seem to be balanced under normal conditions, since *dlk-1(−)* mutants did not show any TRN morphological defects. Perhaps the spatial and temporal expression of DLK-1 determines which of the two opposing functions is engaged. For example, DLK-1 may play inhibitory roles for general neurite extension during the growth phase by reducing MT stability globally in the entire cell and then prevent axon termination at specific sites at a later stage of development. Because *mec-15(−) rpm-1(−)* double mutants exhibited the underextension phenotype, the overall effects of accumulating DLK-1 appear to inhibit neurite growth. Interestingly, DLK-1 also initiates apparently contradictory responses under stress conditions during development and after axonal injury (Tedeschi and Bradke, 2013). For example, after optical nerve injury in mice, DLK triggers the expression of both proapoptotic and regenerative genes, although cell death is the dominant response (Watkins et al., 2013). Future studies are needed to understand how the multifaceted functions of DLK-1 are controlled spatially and temporally to initiate specific signalling. Our work also suggests that different components of the UPS could generate distinct cellular responses by targeting the same protein.

### Inhibitory role of Hsp70/Hsp90 chaperones and co-chaperones in neuronal development

Besides the protective effects of Hsps in cells under stressed conditions, growing evidence suggest that these chaperones also play regulatory roles in normal neurodevelopment (Miller and Fort, 2018). The ATP-dependent Hsp70 and Hsp90 family chaperones are expressed in the nervous system during mouse embryonic and postnatal development, which suggests that they may function in both neurogenesis and late-stage neurodevelopment (Loones et al., 2000). In fact, deletion of the transcription factor HSF1, which activates the transcription of many Hsp genes, led to impaired olfactory neurogenesis and hippocampal spinogenesis and neurogenesis in mice (Uchida et al., 2011), supporting a critical role for Hsps in proper neurodevelopment.

At the subcellular level, Hsc70 (Hsp70 family chaperone) and Hsp90 are localized to the apical dendrites of Purkinje cells and cerebellar neurons from postnatal period into adulthood (D’Souza and Brown, 1998); in the differentiating hippocampal neurons, Hsp90 was associated with the cytoskeleton in branch points and terminal ends (Quinta and Galigniana, 2012). These results suggest that Hsps may regulate neuronal polarization and neurite morphogenesis by modulating cytoskeleton dynamics. Indeed, pharmacological inhibition of Hsp90 disturbed the polarization and axonal extension of cultured hippocampal neurons by inhibiting the PI3K/Akt/GSK3 signaling pathway (Benitez et al., 2014), which is known to regulate neuronal morphogenesis (Kim et al., 2011). The fact that Akt and GSK3 are Hsp90 client proteins (Banz et al., 2009; Sato et al., 2000) indicates that Hsp90 could regulate cell differentiation by maintaining the stability of its client proteins. Thus, given that Hsp90 interacts with hundreds of client proteins, the specific function of Hsp90 in differentiation would depend on the function of its client protein(s) in particular cellular contexts.

Our studies identified one such context, in which Hsp70/Hsp90 chaperones play an unexpected, inhibitory role in regulating MT stability and neurite growth. In *mec-15(−)* mutants, Hsp70/Hsp90 chaperone and co-chaperones contribute to destabilizing MTs, inhibiting neurite growth, suppressing synaptic development, and disrupting sensory functions. Components of this molecular chaperone machinery include HSP-110/Hsp70, HSP-90/Hsp90, cochaperone STI-1/STI1/Hop (which links Hsp70 and Hsp90), PPH-5/PP5 (which interacts with and activates Hsp90 and other cochaperones through dephosphorylation), DAF-41/p23/PTGES3 (which binds to Hsp90 in its ATP-bound conformation and stabilizes the Hsp90-substrate complexes), and CDC-37/Cdc37 (which interacts with Hsp90 and promote the maturation of kinase substrate) (Figure 7). Removing any of these components in *mec-15(−)* mutants rescued the TRN developmental defects and restore normal differentiation. The fact that our experiments identified almost every component in a Hsp90 ATPase cycle (or protein folding cycle) suggests that a complete Hsp90 pathway is involved in negatively regulating TRN differentiation.

**Figure 7.**
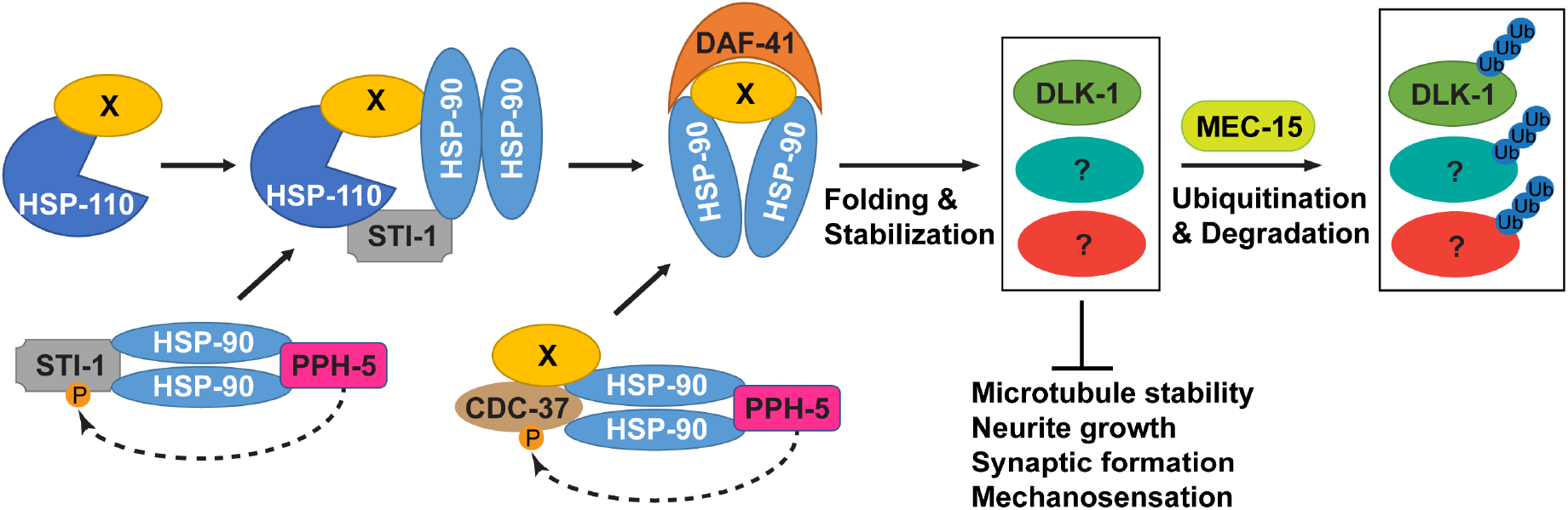
A model for the antagonism between the activities of Hsp70/Hsp90 chaperone machinery and MEC-15 in the regulation of TRN differentiation and functions. HSP-90 receives client protein X from either HSP-110 (a Hsp70 family protein) via a STI-1-mediated physical link or CDC-37 via its binding to kinase clients. Both STI-1 and CDC-37 are activated by PPH-5 through dephosphorylation (dashed line). Those Hsp90 client proteins, including DLK-1 and other unidentified proteins (?), are not only stabilized by the chaperones but also subjected to MEC-15-mediated ubiquitination and degradation. The antagonism between the chaperones and MEC-15 tightly controls the levels of those Hsp90 client proteins, which appear to negatively regulate MT stability, neurite growth, and synaptic development.

At least part of this negative regulation is mediated by the stabilization of DLK-1, which is a conserved Hsp90 client protein in *C. elegans* (this study), Drosophila, and mice (Karney-Grobe et al., 2018). Because the loss of Hsp90 cochaperones fully rescued all developmental defects in *mec-15(−)* mutants, whereas mutations in *dlk-1* only partially suppressed the neurite growth defects, other client proteins stabilized by Hsp90 chaperones may also participate in the inhibition of TRN differentiation. Future studies will focus on identifying such clients.

### The opposing effects of UPS and chaperones in neuronal differentiation

Mutations in the chaperones and cochaperones by themselves did not produce any defects in TRN differentiation, suggesting that the inhibitory function of Hsp70/Hsp90 chaperones and cochaperones does not seem to affect normal development. This lack of effect may be due to Hsp90 clients, such as DLK-1, being normally maintained at a low level by the UPS involving MEC-15, making the effects of chaperone loss difficult to detect. In *mec-15(−)* mutants, when DLK-1 and likely other Hsp90 client proteins accumulate in the TRNs, they are stabilized by the Hsp70/Hsp90 chaperone machinery and exert their functions. In such scenarios, the activity of chaperones in controlling differentiation becomes critical. Thus, by targeting the Hsp90 clients for degradation, UPS curbs the activity of the chaperones in inhibiting cellular differentiation.

These opposing effects of UPS and the chaperones are quite unexpected, since most previous studies indicate that they work synergistically to degrade misfolded proteins. This protein quality control safeguards axon guidance in developing neurons (Wang et al., 2013) and prevents protein aggregation that leads to neurodegeneration (Ciechanover and Kwon, 2017). Our studies, however, provide novel insights into the interaction between UPS and chaperones by suggesting that chaperones also promote the expression and stabilization of proteins that are normally degraded by UPS for fast turnover and low background abundance. This tug-of-war keeps the common substrate of UPS and chaperones at the optimal level, which is crucial for a range of neurodevelopmental processes, including MT formation, neurite growth, synaptic development, and neuronal functions, not only in the TRNs but also in the GABAergic motor neurons. Therefore, we suspect that the balance between the UPS and chaperone activities may be generally important for robust neuronal differentiation.

## Materials and Methods

### Strains and Forward Genetic Screens

*Caenorhabditis elegans* wild-type (N2) and mutant strains were maintained at 20°C as previously described (Brenner, 1974). Alleles used in this study are described in the main text. Most strains were provided by the Caenorhabditis Genetics Center, which is funded by NIH Office of Research Infrastructure Programs (P40 OD010440) or the National BioResource Project of Japan.

For the *mec-7(neo)* suppressor screen, we used TU4879 [*mec-7(u278) X; uIs115 (mec-17p::TagRFP) IV*] as the starter strain, which showed the growth of a long ALM-PN, and ethyl methanesulfonate (EMS) as the mutagen (Brenner, 1974). We isolated 30 mutants that showed very short or no ALM-PN and outcrossed the mutants against TU4879 and identified the phenotype-causing mutations through a combination of whole-genome resequencing, candidate gene sequencing, and complementation tests. *mec-15(u1042)* was isolated from this screen; details about other suppressors will be reported elsewhere.

For the *mec-15(−)* suppressor screen, we used TU5183 [*mec-15(u1042); mec-7(u278); uIs115*] as the starter strain, which had a short or no ALM-PN, and EMS as the mutagen. Through a screen of 29,153 haploid genomes, we isolated 12 mutants that had a prominent ALM-PN with at least five cell body length and identified the phenotype-causing mutations in 8 of the mutants through whole-genome resequencing (Table S1). Deletion alleles *sti-1(ok3354)* and *pph-5(ok3498)* and frameshift-causing allele *dlk-1(ju476)* were used as the reference alleles.

### Candidate RNAi Screen

To test Skp and Cullin homologs for the suppression of *mec-7(u278)*, we constructed TU6023 *[eri-1(mg366ts) uIs115 IV; mec-7(u278) lin-15b(n744) X]* for enhanced RNAi effects and performed feeding RNAi using bacteria colonies from the Ahringer library (Fraser et al., 2000). Briefly, we seeded bacteria expressing dsRNA on NGM agar plates containing 6 mM IPTG and 100 μg/ml ampicillin. One day later, eggs from TU6023 animals were placed onto these plates, hatched, and grown to adults at 20°C. The F1 progeny of these worms were examined for the growth of ALM-PN at the day one adult stage. Genes whose knockdown shortened the ALM-PN length to below five cell body lengths were regarded as positive.

To test candidate genes for the suppression of *mec-15(−)* mutation, we constructed TU6092 *[mec-15(u75); eri-1(mg366) uIs115; mec-7(u278) lin-15B(n744)]* and performed feeding RNAi for the genes indicated in Table S2. Genes whose knockdown restored the ALM-PN length to above five cell body lengths were regarded as positive; positives from the screen were confirmed using deletion alleles, including *hsp-90(ok1333)*, *hsp-110(gk533)*, and *daf-41(ok3052)*.

### Constructs and transgenes

For *mec-15* reporter and expression constructs, a *mec-15::GFP* translational reporter (TU#2278) was made by cloning a 3.2 kb promoter upstream of the translational start and the entire *mec-15* coding sequence into the pPD95.75 backbone and was used to generate an integrated transgene *unkIs1[mec-15::GFP]*. *mec-18p::mec-15* (TU#891) was generated previously by inserting a 0.5-kb *mec-18* promoter and the *mec-15* coding sequence with a 1.6 kb 3’UTR into the pBlueScript II backbone (Bounoutas et al., 2009b). *mec-18p::mec-15(Δ9-50)* (TU#2100), *mec-18p::mec-15(Δ9-84)* (TU#2101), and *mec-18p::mec-15::HA* (CGZ#84) were generated via site-directed mutagenesis using TU#891 as a template.

For *pph-5* and *sti-1* reporter and expression constructs, the *pph-5* transcriptional reporter transgene *avIs90[pph-5p::mCherry::H2B::pph-5 3’UTR; unc-119(+)]* was generated by Richie et al. (2011). For the *pph-5* translational reporter, we amplified a 2.3-kb promoter, the cDNA of *pph-5* (isoform a), and a 481-bp *pph-5* 3’UTR and assembled the PCR products with amplified pUC57 backbone via HiFi assembly (New England Biolabs) to generate the vector *pph-5p::pph-5(cDNA)::pph-5 3’UTR* (TU#2257). The GFP coding sequence was then subcloned from pPD95.75 and inserted before the stop codon of *pph-5* to generate *pph-5p::pph-5(cDNA)::GFP::pph-5 3’UTR* (TU#2267) via HiFi assembly. The *sti-1p::GFP::unc-54 3’UTR* (TU#2202) transcriptional reporter was generated through the Gateway system (Life Technologies) by cloning a 1045-bp promoter into pDONR-P4-P1r and then performing LR reactions with pENTR-GFP, pENTR-unc-54-3’ UTR, and destination vector pDEST-R4-R3. We then generated *sti-1p::sti-1::sti-1 3’UTR* vector (TU#2264) by amplifying the backbone plus the *sti-1* promoter from TU#2202 and the *sti-1* genomic coding sequence plus a 607-bp 3’UTR from genomic DNA and assembled them via HiFi assembly. GFP-coding sequence was then inserted before the *sti-1* stop codon to generate *sti-1p::sti-1::GFP::sti-1 3’UTR* (TU#2280).

For TRN-specific expression or RNAi constructs, *mec-17p::pph-5(cDNA)::pph-5 3’UTR* (TU#2266) was generated by swapping the *pph-5* promoter in TU# 2257 with a 2.9-kb *mec-17* promoter. *mec-17p::sti-1::unc-54 3’UTR* (TU#2166), *mec-17p::daf-41::FLAG::unc-54 3’UTR* (CGZ#92), *mec-17p::FLAG::dlk-1a::unc-54 3’UTR* (CGZ#78) were generated by cloning *mec-17* promoter and *sti-1*, *daf-41*, and *dlk-1a* genomic coding sequences into the Gateway vectors and then performing the LR reactions with pENTR-unc-54-3’ UTR and pDEST-R4-R3. GFP was inserted into CGZ#78 after the start codon of *dlk-1* to make *mec-17p::GFP::dlk-1a::unc-54 3’UTR* (CGZ#71). Various *mec-17p::sti-1* mutant constructs were generated by site-directed mutagenesis using TU#2166 as a template. For TRN-specific RNAi of *uba-1*, *hsf-1*, and *cdc-37*, we amplified the sense and antisense sequences of their genomic coding region (without start and stop codons), and inserted them downstream of a *mec-17* promoter in pDEST-R4-R3 backbones; the resulting constructs *mec-17p::uba-1-sense* (TU#2286), *mec-17p::uba-1-antisense* (TU#2287), *mec-17p::hsf-1-sense* (CGZ#97), *mec-17p::hsf-1-antisense* (CGZ#98), *mec-17p::cdc-37-sense* (CGZ#82), and *mec-17p::cdc-37-antisense* (CGZ#83) were injected in pairs into *mec-7(u278)* or *mec-15; mec-7* animals for RNA interference. As negative controls for RNAi, we injected *mec-17p::gfp-sense* (CGZ#167) and *mec-17p::gfp-antisense* (CGZ#168) into corresponding animals to express dsRNA against GFP.

To generate transgenic animals, we injected DNA constructs (5 ng/μl for each expression vector) into the animals to establish stable lines carrying an extrachromosomal array; at least three independent lines were tested. In some cases, we integrated the arrays into the genome through standard γ-irradiation (Mello and Fire, 1995) to generate integrated transgenes, including *unkIs1[mec-15::GFP]* and *unkIs2[mec-17p::GFP::dlk-1]*. Those integrated lines were outcrossed for at least 5 times before being examined.

Transgenes *uIs31[mec-17p::GFP] III*, *uIs115[mec-17p::RFP] IV*, and *uIs134[mec-17p::RFP] V* were used to visualize TRN morphology (Zheng et al., 2015). *jsIs821[mec-7p::GFP::RAB-3]* was used to assess synaptic vesicle localization (Bounoutas et al., 2009b). *sEx10796[daf-41p::GFP]* was used to examine *daf-41* expression pattern. *juIs1[unc-25p::SNB-1-GFP]* was used to examine synaptic puncta in the GABAergic motor neurons.

### Electron microscopy

We fixed TU4065 [*uIs115(mec-17p::TagRFP)*], TU5598 [*mec-15(u1042); uIs115*], TU5579 [*pph-5(ok3498); mec-15(u1042); uIs115*], and TU5580 [*sti-1(ok3354); mec-15(u1042); uIs115*] adults grown at 20°C using either a chemical immersion (Hall, 1995) or a modified high-pressure freezing/freeze substitution (HPF/FS) protocol (Zheng et al., 2017). Chemical immersion was performed using 1% glutaraldehyde + 1% tannic acid in 0.1 M phosphate buffer (pH7.0); and HPF/FS was done by a first fixation in 0.5% glutaraldehyde + 0.2% tannic acid at −90°C for 90 hours and then at −60°C for 4 hours, followed by a second fixation in 2% osmium tetroxide + 0.2% uranyl acetate at −60°C for 16 hours and then at −30°C for 16 hours.

Eighty-nanometer transverse sections were collected at multiple positions along both the ALM anterior neurite and the PLM anterior neurites and poststained with uranium acetate and lead citrate. JEOL JEM-1400Plus TEM with a Galan Orius SC1000B bottom-mount digital camera or a Philips CM10 electron microscope with an attached Morada digital camera (Olympus) was used to acquire the images. For each strain, we examined sections from at least ten animals. For each animal, we collected sections from at least five different positions along the neurite and counted the number of MTs in a cross-section, measured the MT diameter, and counted the number of protofilaments in individual MTs, of which clear substructure were observed. Data from ALML and ALMR were combined and PLML and PLMR combined. Number of data points for each measurement is indicated in Figure 4B and 4D.

### Yeast two-hybrid assay and western blot

Yeast media and plates were prepared according to recipes from Clontech and yeasts were grown at 30°C. The yeast strain PJ69-4a used for the two-hybrid assays contains GAL1-HIS3, GAL2-ADE2 and GAL7-lacZ reporters (Zheng et al., 2018). Vectors pGAD424 and pGBT9 (Clontech) were used to express proteins fused to the yeast-activating domain (AD) and -binding domain (BD), respectively. cDNA fragments of *mec-15*, *skr-1*, *hsp-90*, and *dlk-1a* were cloned into the two-hybrid vectors using HiFi Assembly (NEB).

Combinations of the AD or BD vectors were co-transformed into yeast using the Frozen-EZ II kit from Zymo Research and using empty vectors as negative controls. Growth assays were performed by growing individual colonies overnight in selective media lacking tryptophan and leucine and then diluting the cultures to OD600 = 0.5; a series of further 1:10 dilutions were made, and 10 μl of the diluted culture were spotted onto plates lacking histidine to test the expression of the HIS3 reporter. Plates were imaged after 2 days of growth.

For western blot, transgenic animals were collected and lysed with the RIPA buffer [25 mM Tris-HCl, pH 7.5; 100 mM NaCl, 1 mM EDTA, 0.5% NP-40, and freshly added protease inhibitors] and sonicated for 15 cycles of 1 second on and 15 seconds off. The protein lysate was then separated by SDS-PAGE and blotted with anti-HA (Cat# HT301-02, TransGen Biotech, Beijing) and anti-FLAG (Cat# F1804, Sigma) primary antibodies and HRP-conjugated secondary antibodies (Cat# A9044, Sigma).

### Fluorescent imaging, aldicarb assay, phenotype scoring and statistical analysis

Fluorescent imaging was done either on a Zeiss Axio Observer Z1 inverted microscope equipped with a CoolSNAP HQ2-FW camera (Photometrics) or a Leica DMi8 inverted microscope equipped with a Leica DFC7000 GT monochrome camera. Measurements were made using either the Zeiss AxioVision (SE64 Rel. 4.9.1) or the Leica Application Suite X (3.7.0.20979) software.

We measured the length of TRN neurites in at least 20 larvae or young adults grown at 20°C, except where otherwise stated. Normalized neurite length was calculated by dividing the neurite length by the diameter of the cell body along A-P axis. Defects in PLM-AN growth were assessed by measuring the distance from the PLM-AN terminals to the center of the vulva. Positive numbers mean PLM-AN terminals are more anterior than the vulva; negative numbers mean PLM-AN did not reach the vulva; at least 30 cells were examined for each strain. All bar graphs show mean ± SD.

Fluorescence intensity of the TagRFP expressed from the *uIs115[mec-17p::TagRFP]* or the GFP expression from *unkIs2[mec-17p::GFP::dlk-1]* transgene in the PLM cell body was used to measure TRN protein levels; intensity was measured by first acquiring images using the identical exposure time and laser power and then manually subtracting the background from the fluorescent signal using ImageJ. At least 30 cells were imaged and analyzed for each strain.

For the assay of sensitivity to aldicarb, an acetylcholinesterase inhibitor, 20 day-one adult animals were transferred onto NGM plates containing 1 mM aldicarb (Sigma) without food. The number of animals that were paralyzed was recorded every 10 minutes, starting at 30 minutes after the transfer. Genotypes were blind to the experimenter, and the assay was repeated three times.

For statistical analysis, analysis of variance and the post hoc Dunnett’s test (comparison of each of many treatments with a single control) or Tukey–Kramer test (studentized range test for all pair-wise comparison) were performed using the GraphPad Prism (version 8.00 for Mac, GraphPad Software, La Jolla, CA; www.graphpad.com) to identify significant difference among samples. Student’s t test was used to find significant difference between two samples in paired comparisons. Single and double asterisks indicate p < 0.05 and p < 0.01, respectively.

## Acknowledgements

We thank Natalie Yvonne Sayegh for helping with the *mec-7* suppressor screen, Andy Golden for sharing strains carrying *avIs90*, and the members of the Chalfie lab for constructive comments on the manuscript. This work is supported by start-up funds from the University of Hong Kong to C.Z. and grants from the Research Grant Council of Hong Kong [ECS27104219 to C.Z.] and the National Institutes of Health of United States [GM30997 and GM122522 to M.C.; OD010943 to D.H.H.]. The electron microscope JEOL 1400Plus TEM used in this study was acquired through a NIH Shared Instrumentation Grant (1S10OD016214-01A1) at Albert Einstein College of Medicine.

## Author Contributions

Conceptualization, C.Z. and M.C.; Methodology, C.Z., E.A., K.C.Q.N, D.H.H., and M.C.; Investigation, C.Z., E.A., H.M.T.L, S.L.J.J, and K.C.Q.N; Writing – Original draft and revision, C.Z. and M.C.; Funding acquisition, C.Z., D.H.H., and M.C.; Resource, D.H.H.; Supervision, C.Z. and M.C.

