## Supplementary material for "Opposing effects of an F-box protein and the HSP90 chaperone network on microtubule stability and neurite growth in *Caenorhabditis elegans*": Table S1-S2; Figure S1-S7

**Supplemental Materials**

**Supplemental Results - An unexcepted maternal effect of chaperone and cochaperone mutants**

**Table S1. Summary of the mutants isolated from the *mec-15(-)* suppressor screen.**

**Table S2. Results of the candidate RNAi screen for *mec-15(-)* suppressors.**

**Figure S1. F-box protein MEC-15 is required for TRN neurite development.**

**Figure S2. Mutations or knockdown of Hsp90 cochaperones suppress the phenotype of *mec-15* mutants.**

**Figure S3. Hsp90 cochaperones are expressed in the TRNs and function cell-autonomously.**

**Figure S4. Maternal effects of Hsp70/Hsp90 chaperone and cochaperone mutants.**

**Figure S5. Single mutants of Hsp70/90 chaperones and cochaperones showed normal TRN morphologies.**

**Figure S6. Mutations in *dlk-1* fail to rescue the defects in TRN synaptic development and sensory function of *mec-15* mutants.**

**Figure S7. MEC-15 downregulates DLK-1 and is epistatic to RPM-1.**

### Supplemental Results

#### An unexcepted maternal effect of chaperone and cochaperone mutants

Mutants of chaperones and cochaperones had unexpected, strong maternal effects in suppressing the Mec-15 phenotype. For example, although paternally derived *sti-1/+; mec-15; mec-7* heterozygotes had short or no ALM-PN similar to *mec-15; mec-7* animals, heterozygotes produced by *sti-1; mec-15*; *mec-7* homozygous mothers showed long ALM-PN (Figure S4A). This result suggested that despite the presence of zygotic *sti-1(+)*, the lack of maternally deposited STI-1 mRNAs or proteins into the oocytes was sufficient to suppress the defects caused by *mec-15(lf)* mutations. Consistent with the maternal effect, *sti-1; mec-15*; *mec-7* homozygotes produced by the *sti-1/+; mec-15; mec-7* heterozygous mother had very short ALM-PN, whereas homozygotes derived from homozygous mother had long ALM-PN (Figure S4C). Similar maternal effects were observed with the *sti-1; mec-15* double mutants in the absence of the *mec-7(u278 neo)* background, for the suppression of PLM neurite growth defects (Figure S4B).

To reconcile this maternal effect with the cell-autonomous rescue of *sti-1* mutants, we hypothesize that the expression of endogenous *sti-1(+)* gene in TRNs may be at a low level or have a late onset, which causes a dependency on the maternal contributed protein. Overexpression of *sti-1(+)* from the *mec-17* promoter, a strong TRN-specific promoter that is activated shortly after the generation of TRNs, may overcome the lack of maternally deposited STI-1.

Mutations in *pph-5* and *daf-41/p23* had similar maternal effects in either the *mec-15; mec-7* double mutants or the *mec-15* single mutants (Figure S4A-C). Although *hsp-110(gk533)* homozygous deletion mutants arrested at early larval stages, a few in each generation escaped, became vulvaless adults, and produced about ~20 progeny. Those *hsp-110; mec-15; mec-7* triple mutants derived from homozygous mothers had a longer ALM-PN than the ones derived from the *hsp-110/nT2; mec-15; mec-7* heterozygous mothers; *hsp-110; mec-15* double mutants produced by homozygous mothers also had a longer PLM-PN than the ones produced by the heterozygous mothers (Figure S4D-E). Thus, mutations in *hsp-110* (Hsp70) also showed maternal effects. We could not test the *hsp-90* mutants for maternal effects because no escapers were found.

Overall, our results suggest that maternally deposited chaperone and cochaperone mRNAs or proteins have long-lasting effects on the regulation of neurite morphogenesis in terminally differentiated TRN neurons. This observation is quite surprising, given that the generation of those neurons occurs more than 7 hours after fertilization at 20°C and requires 10 (for ALMs) or 12 (for PLMs) rounds of embryonic cell divisions from the zygote (Sulston et al., 1983) and that neurite extension continuously occurs for three days in post-embryonic development after hatching.

| Gene name | Allele name | Molecular lesion | Recessivity | Maternal effect |
|---|---|---|---|---|
| *sti-1* | *u1071* | R253* | recessive | Yes |
| *pph-5* | *u1072* | C381Y | recessive | Yes |
| *pph-5* | *u1073* | P448L | recessive | Yes |
| *pph-5* | *u1074* | Q325* | recessive | Yes |
| *pph-5* | *u1075* | Q192* | recessive | Yes |
| *dlk-1* | *u1105* | V844I | recessive | Yes |
| *dlk-1* | *u1138* | W394* | recessive | Yes |
| *maph-1.3* | *u1107* | E794K | dominant | N/A |
| ? | *u1108* | | dominant | N/A |
| ? | *u1106* | | semi-dominant | N/A |
| ? | *u1142* | | semi-dominant | N/A |
| ? | *u1144* | | semi-dominant | N/A |

**Table S1. Summary of the mutants isolated from the *mec-15(-)* suppressor screen.** These alleles led to the production of a long ALM-PN in *mec-15(u1042); mec-7(u278); uIs115[mec-17p::TagRFP]* animals. * represents a stop codon; ? means the mutant allele was not mapped.

| Gene name | Transcript name | Description | Long ALM-PN |
|---|---|---|---|
| *pph-5* | Y39B6A.2 | Protein Phosphatase 5 | Positive |
| *sti-1* | R09E12.3 | Hsp90 co-chaperone; Hop/STI1 | Positive |
| *hsp-1* | F26D10.3 | Hsp70 family | Negative |
| *hsp-110* | C30C11.4 | HSPA4; Hsp70 family | Positive |
| *hsp-3* | C15H9.6 | HSPA5; Hsp70 family | Negative |
| *hsp-4* | F43E2.8 | HSPA5; Hsp70 family | Negative |
| *hsp-6* | C37H5.8 | HSPA9; Hsp70 family | Negative |
| *stc-1* | F54C9.2 | HSPA13; Hsp70 family | Negative |
| *hsp-70* | C12C8.1 | Hsp70 family | Negative |
| *F44E5.4* | F44E5.4 | Hsp70 family | Negative |
| *F44E5.5* | F44E5.5 | Hsp70 family | Negative |
| *T14G8.3* | T14G8.3 | HYOU1; Hsp70 family | Negative |
| *hsp-75* | R151.7 | Hsp90-related protein; Hsp75/TRAP1 | Negative |
| *enpl-1* | T05E11.3 | Molecular Chaperone; GRP94/GP96 | Negative |
| *chn-1* | T09B4.10 | CHIP; Hsp70-interacting protein | Negative |
| *daf-41* | ZC395.10 | Hsp90 co-chaperone; p23 | Positive |
| *aha-1* | C25A1.11 | Hsp90 co-chaperone; Aha1 | Negative |
| *aipr-1* | C56C10.10 | Hsp90 co-chaperone; XAP2 | Negative |
| *bag-1* | F57B10.11 | Hsp90 co-chaperone; Bag-1 | Negative |
| *fkb-6* | F31D4.3 | FK506-Binding protein; Hsp90-interacting protein | Negative |
| *cdc-37* | W08F4.8 | Hsp90 co-chaperone; cdc37 | Negative |
| *trak-1* | T27A3.1 | HAP1; Hsp90-interacting protein | Negative |
| *tomm-70* | ZK370.8 | TPR repeat-containing mitochondrial protein | Negative |
| *sgt-1* | R05F9.10 | PPH-5 homolog; TRP repeat-containing protein | Negative |
| *cct-1* | T05C12.7 | Chaperonin Containing TCP-1 | Negative |
| *hsb-1* | K08E7.2 | Heat shock factor binding protein | Negative |
| *lit-1* | W06F12.1 | Serine/threonine-protein kinase NLK | Negative |

**Table S2. Results of the candidate RNAi screen for *mec-15(-)* suppressors.** *mec-15(u75); eri-1(mg366) uIs115; mec-7(u278) lin-15B(n744)* animals were fed with RNAi bacteria. If 10% of the treated animals had ALM-PN longer than five cell body length, the treatment was considered to produce positive results.

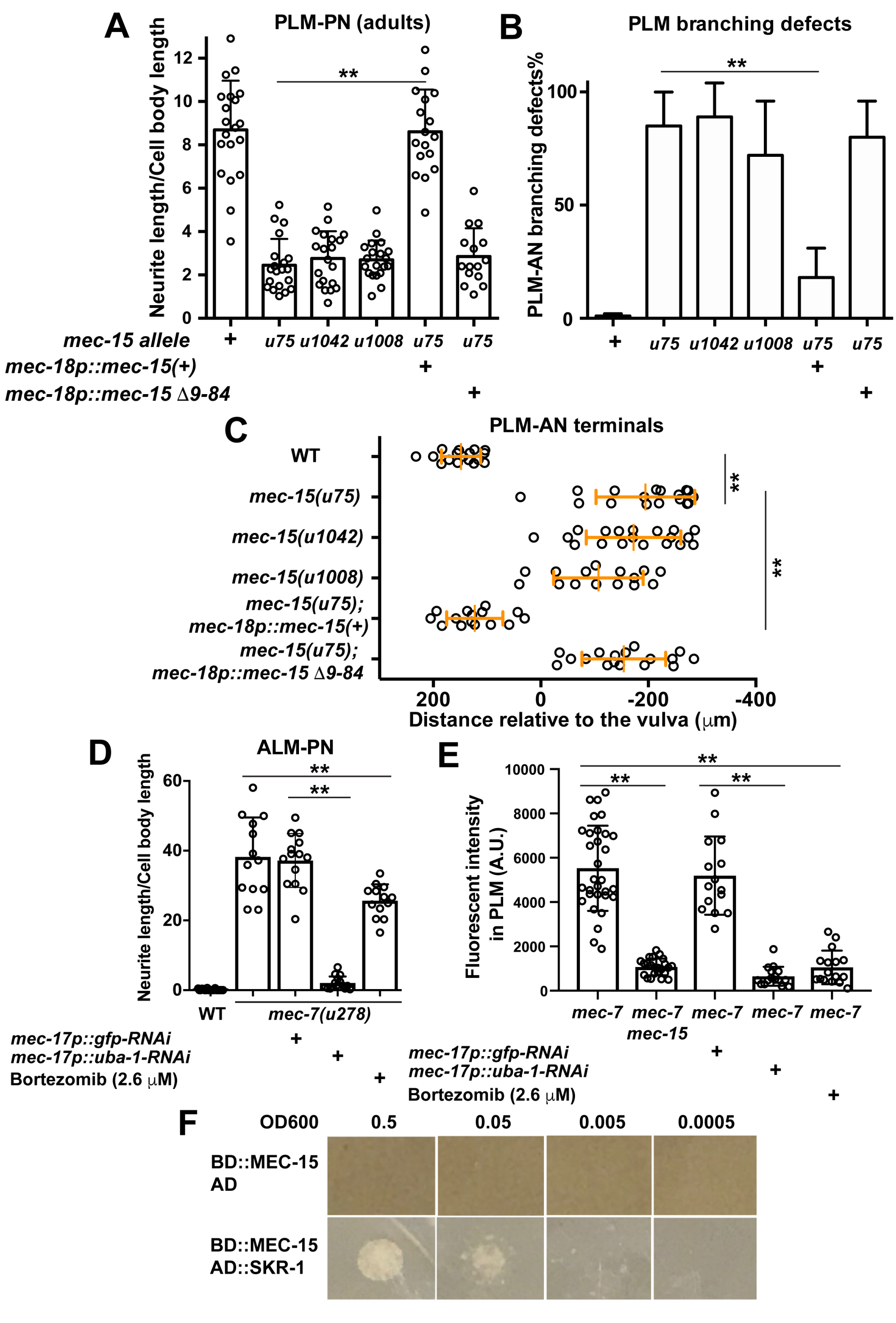


**Figure S1. F-box protein MEC-15 is required for TRN neurite development.** (A) The length of PLM-PN in animals carrying various *mec-15 lf* alleles and *mec-15(u75)* mutants carrying the transgene expressing either wild-type MEC-15 (+) or MEC-15 Δ9-84 truncates from the TRN-specific *mec-18* promoter. (B) The percentage of PLM cells with PLM-AN branching defects in various strains. N > 40. (C) The distance of PLM-AN terminals to the vulva in the indicated strains. The distance is positive if PLM-AN grew pass the vulva towards the anterior and negative if PLM-AN did not reach the vulva.

(D) The length of ALM-PN in wild type (WT), *mec-7(u278)* mutants, and *mec-7* animals carrying the transgene expressing dsRNA against *uba-1* from the TRN-specific *mec-17* promoter, as well as in *mec-7* animals grown on plates containing 2.6 μM bortezomib. (E) TagRFP fluorescent intensity in PLM neurons of *mec-7(u278)* mutants, *mec-7 mec-15(u1042)* double mutants, and *mec-7* animals with *uba-1* knockdown or bortezomib treatment. (F) Yeast two-hybrid assays for the interaction between MEC-15 and SKR-1. AD means the GAL4 activation domain and BD means the GAL4 DNA-binding domain.

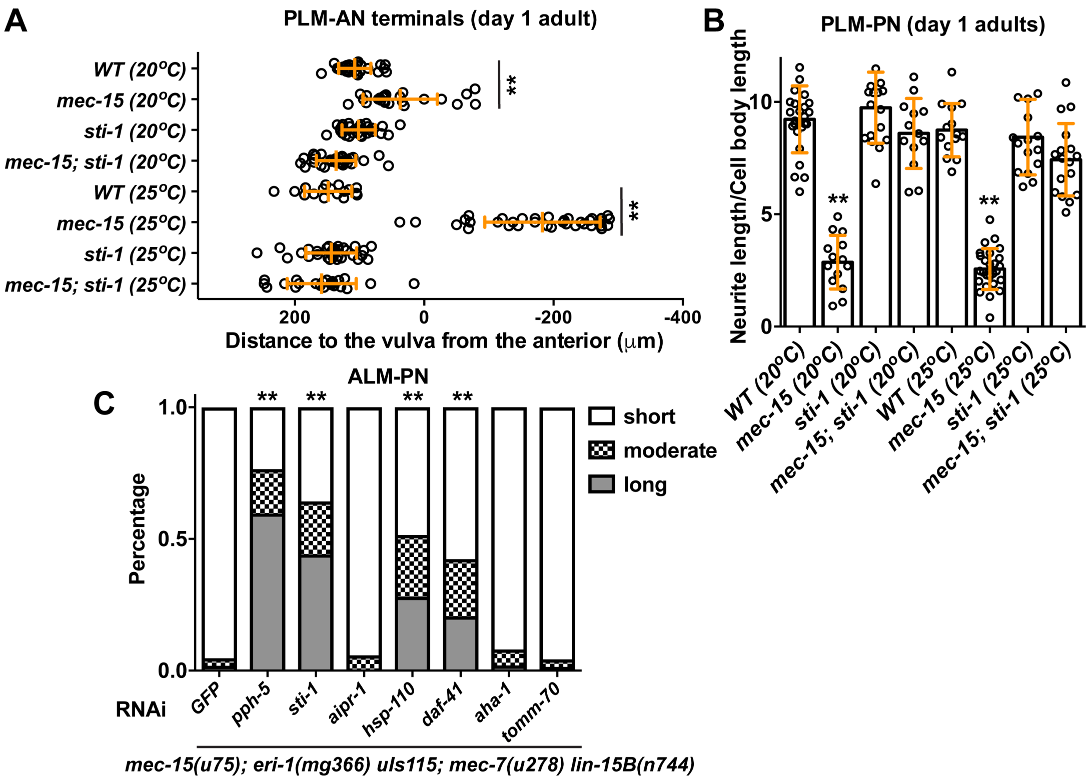


**Figure S2. Mutations or knockdown of Hsp90 cochaperones suppress the phenotype of *mec-15* mutants.** (A) The distance of PLM-AN terminals to the vulva in wild type (WT) *mec-15(u1042)*, *sti-1(ok3354)* and *mec-15; sti-1* animals at both 20 and 25 °C. (B) The length of PLM-PN in *mec-15(u1042)*, *sti-1(ok3354)* and *mec-15; sti-1* animals at both 20 and 25 °C. Double asterisks indicate statistical significance ($p < 0.01$) for the difference between *mec-15* mutants and the wild type animals. (C) The percentage of ALM cells with short (< 2 cell body length), moderate (2-5 cell body length), or long (> 5 cell body length) ALM-PN in *mec-15; eri-1 uIs115[mec-17p::TagRFP]; mec-7 lin-15B;* animals fed with bacteria that express dsRNA against the genes indicated. Double asterisks indicate statistical significance ($p < 0.01$) for the difference between the treatment and the RNAi against GFP in a Chi-square test.

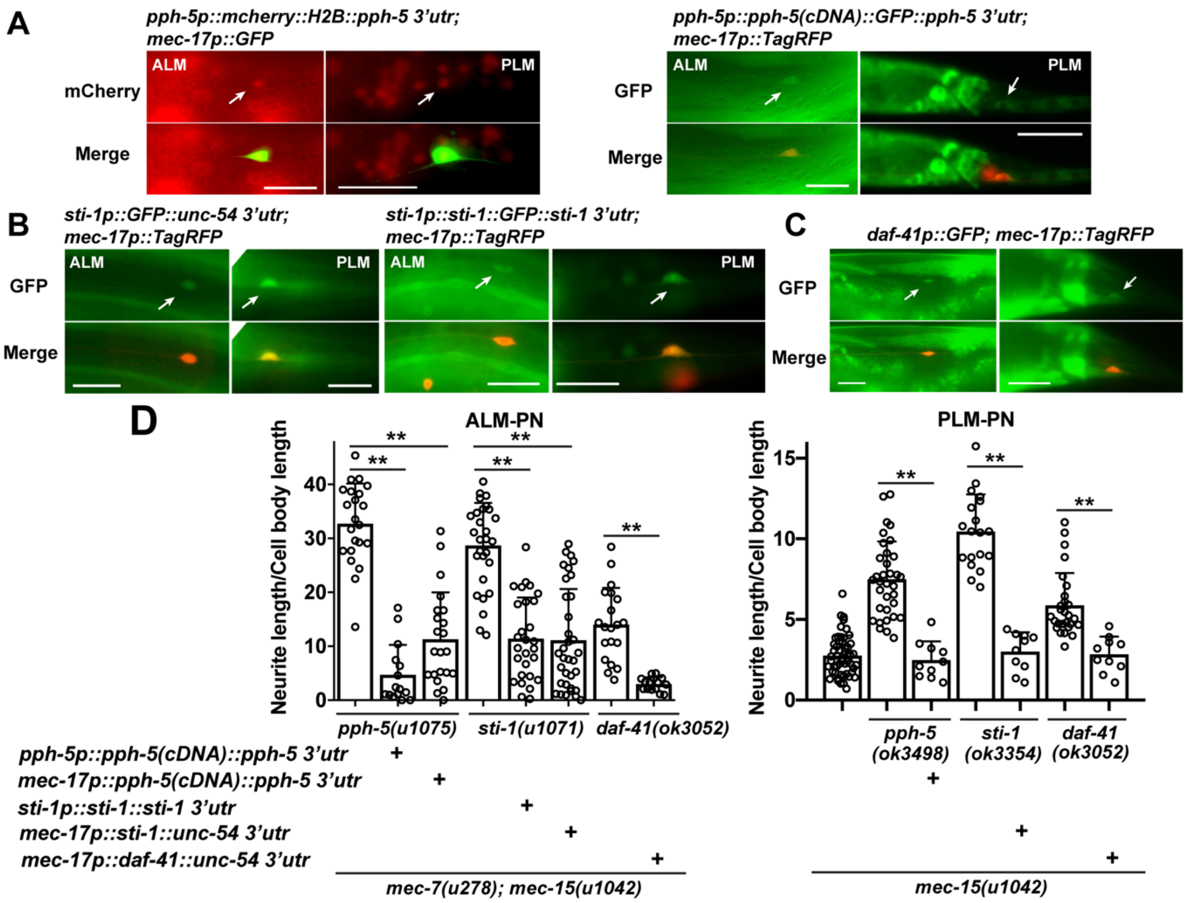


**Figure S3. Hsp90 cochaperones are expressed in the TRNs and function cell-autonomously.** (A) The expression of *pph-5* transcriptional reporter *avIs90[pph-5p::mCherry::H2B::pph-5 3'utr; unc-119(+)]* (Richie et al., 2011) and translational reporter *unkEx36[pph-5p::pph-5_cDNA::GFP::pph-5 3'utr; unc-119(+)]* in TRNs. (B) The expression of *sti-1* transcriptional reporter *unkEx52[sti-1p::GFP::unc-54 3'utr]* and translational reporter *unkEx24[sti-1p::sti-1::GFP::sti-1 3'utr; unc-119(+)]* in TRNs. (C) The expression of *daf-41* transcriptional reporter *sEx10796[daf-41p::GFP]* in TRNs. Arrows point to the cell bodies of ALM and PLM. Scale bar = 20 μm. (D) The length of ALM-PN in the triple mutants that expressed the rescuing transgenes and the length of PLM-PN in the double mutants that expressed the transgenes. Double asterisks indicate statistical significance ($p < 0.01$).

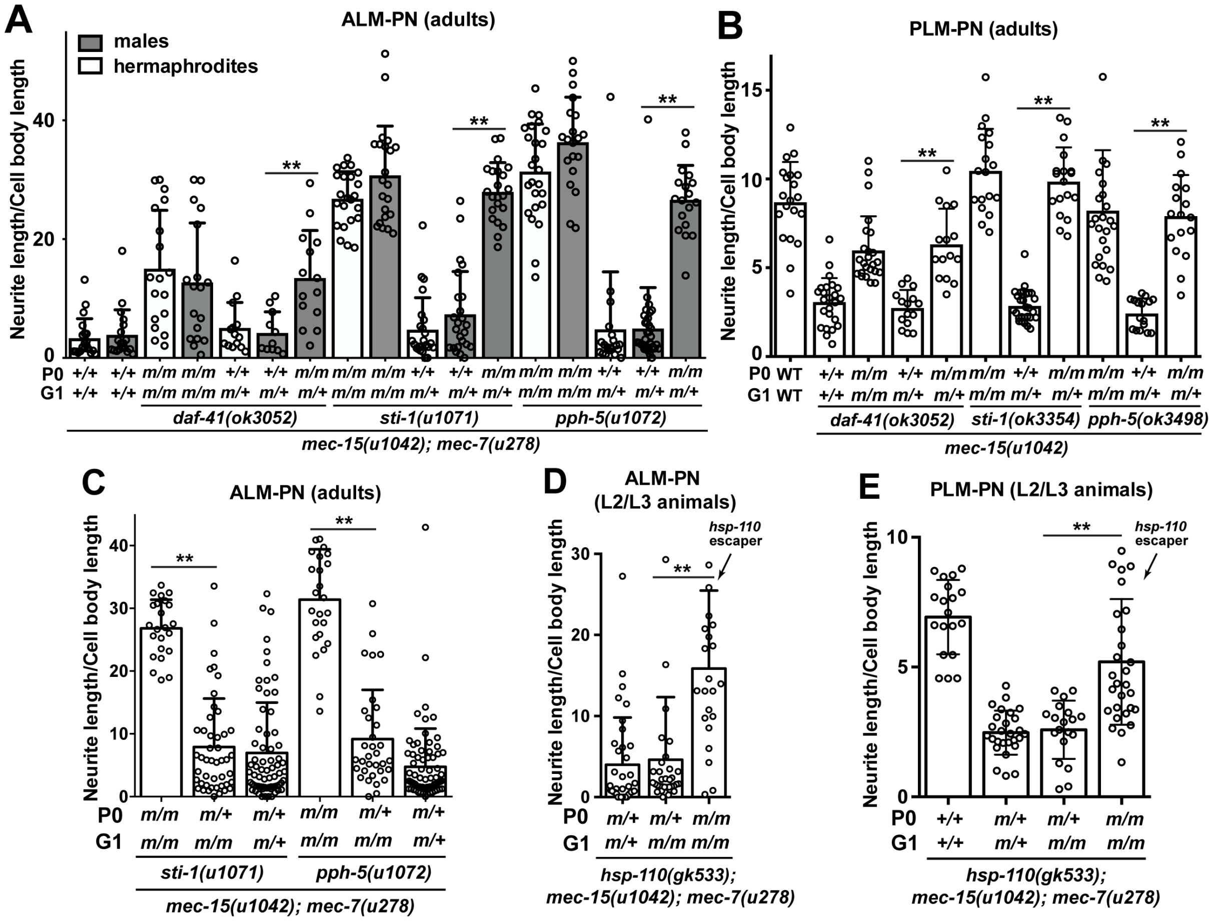


**Figure S4. Maternal effects of Hsp70/Hsp90 chaperone and cochaperone mutants.** (A) The length of ALM-PN of the heterozygous cross-progeny between *mec-15; mec-7* double mutants and their triple mutants with *daf-41*, *sti-1*, or *pph-5 lf* mutations. P0 indicates the genotype of the mother of the animals (G1 for generation 1) that were examined; "m" represents the mutant allele and "+" the wild-type allele. Crosses to generate heterozygous progeny were repeated three times. (B) The length of PLM-PN of the heterozygous cross-progeny between *mec-15* single mutants and the *mec-15* double mutants with *daf-41*, *sti-1*, or *pph-5 lf* mutations. (C-D) The length of ALM-PN of the self-progeny of *sti-1/+; mec-15; mec-7* and *pph-5/+; mec-15; mec-7* animals and the progeny of *hsp-110/+; mec-15; mec-7* hermaphrodites and the *hsp-110; mec-15; mec-7* escapers. (E) The length of PLM-PN of the self-progeny of *hsp-110/+; mec-15;* hermaphrodites and the *hsp-110; mec-15* escapers. *hsp-110* homozygotes mostly arrested at larval stages.

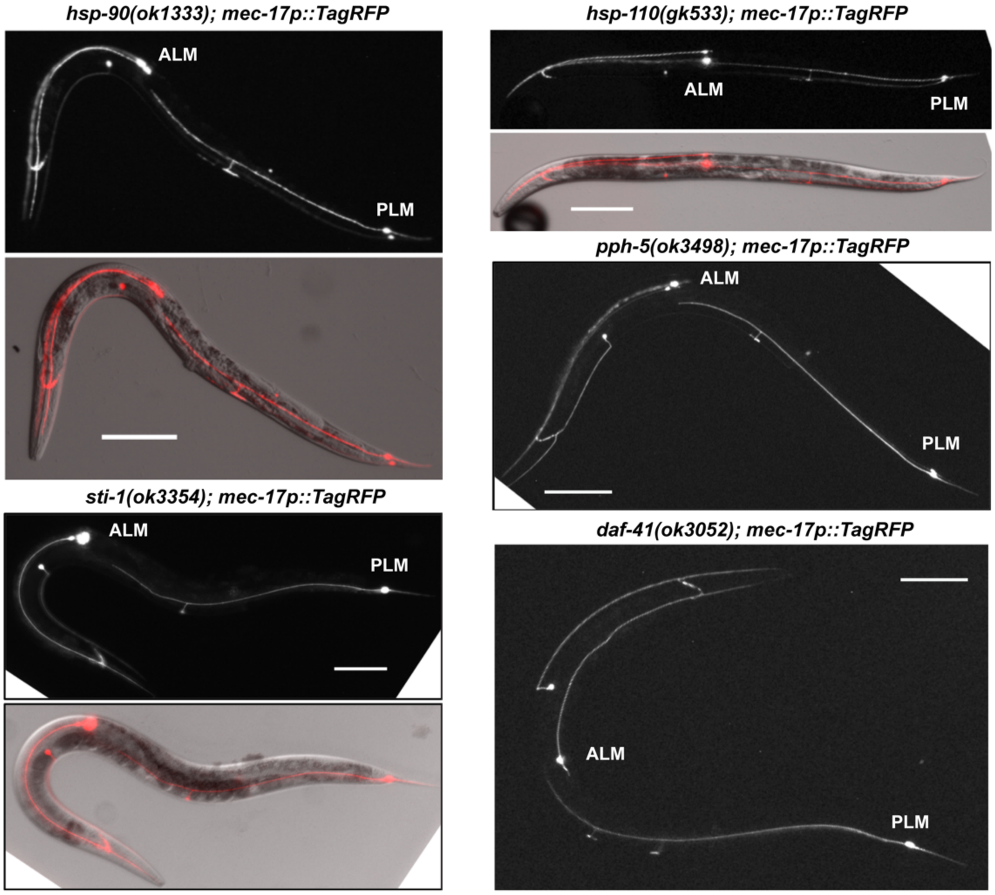


**Figure S5. Single mutants of Hsp70/90 chaperones and cochaperones showed normal TRN morphologies.** TRN morphologies were visualized by the *mec-17p::TagRFP* transgene in *hsp-90*, *hsp-110*, *sti-1*, *pph-5*, and *daf-41* mutants. *hsp-90* and *hsp-110* homozygotes were the progeny of the balanced heterozygotes *hsp-90/nT1* and *hsp-110/hT2*, respectively. Scale bars = 100 µm.

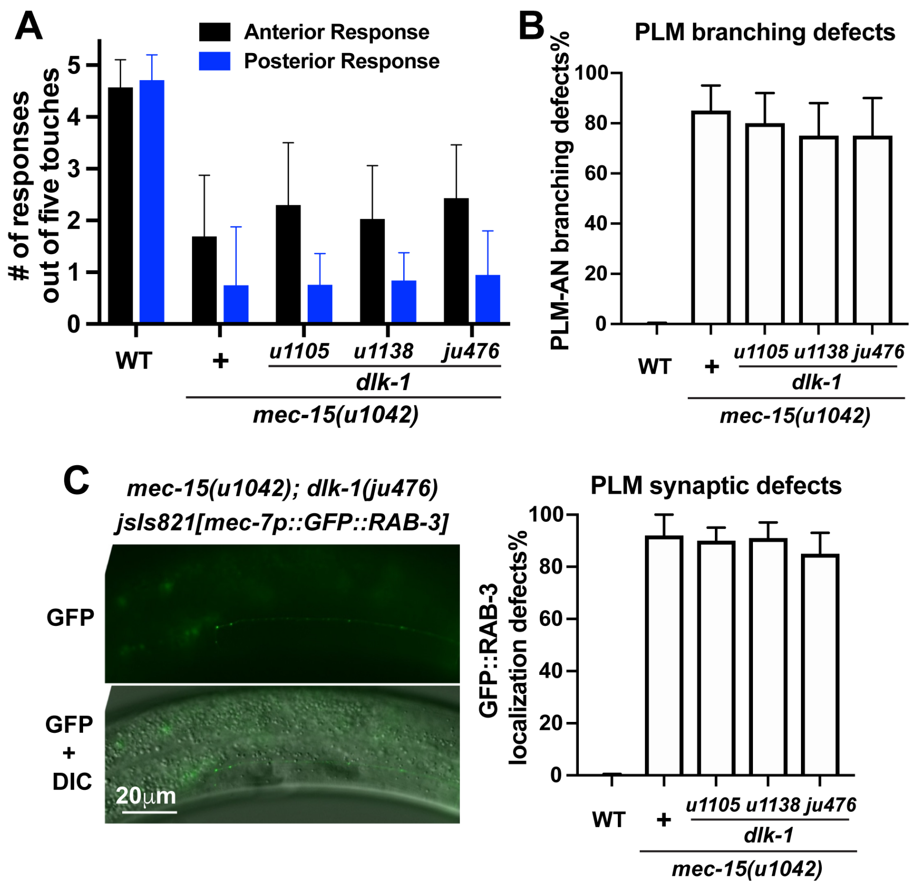


**Figure S6. Mutations in *dlk-1* fail to rescue the defects in TRN synaptic development and sensory function of *mec-15* mutants.** (A) The number of gentle touch response out of five stimuli in *mec-15* and *mec-15 dlk-1* mutants. (B) The percentages of PLM cells with PLM-AN branching defects in *mec-15* and *mec-15 dlk-1* mutants. (C) Defects of synaptic vesicle (GFP::RAB-3) localization in *mec-15 dlk-1* mutants. The percentage of PLM cells with abnormal GFP::RAB-3 localization in *mec-15* and *mec-15 dlk-1* mutants.

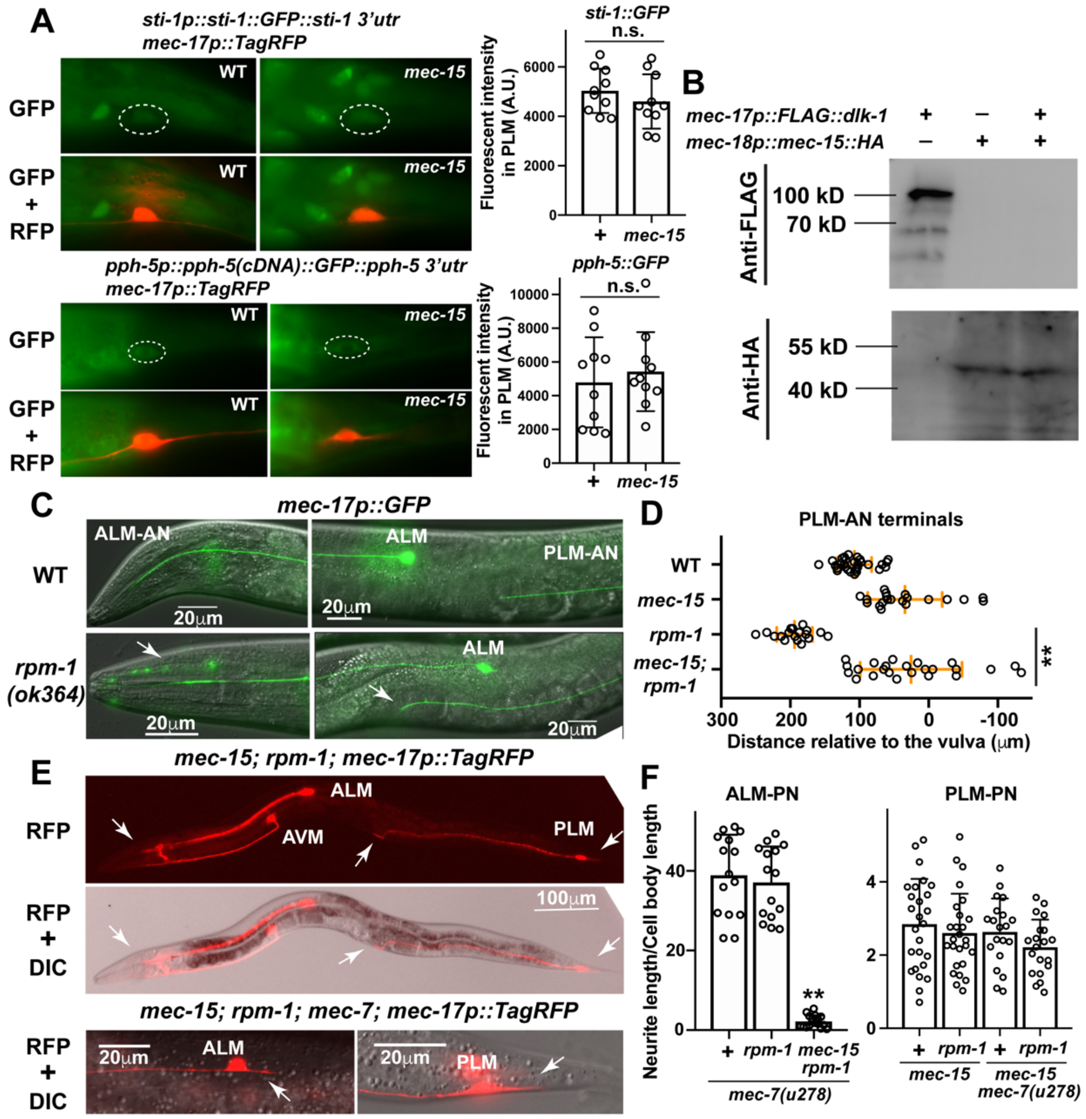


**Figure S7. MEC-15 downregulates DLK-1 and is epistatic to RPM-1.** (A) Fluorescence of STI-1::GFP and PPH-5::GFP in PLMs of the wild-type and *mec-15(u1042)* animals. Dashed circles enclosed the PLM cell bodies. Quantification of the GFP fluorescent intensity in PLM cells are shown. (B) Western blot with anti-FLAG or anti-HA antibodies with animals carrying transgenes expressing either FLAG::DLK-1 or MEC-15::HA or both in TRNs. (C) Overextension of ALM-AN and PLM-AN in *rpm-1(ok364)* mutants. (D) The distance of PLM-AN terminals to the vulva in *mec-15(u1042)*, *rpm-1(ok364)*, and *mec-15; rpm-1* mutant animals. (E) The shortening of ALM-AN, PLM-AN, and PLM-PN in *mec-15; rpm-1* mutant and the shortening of ALM-PN and PLM-PN in *mec-15; rpm-1; mec-7* animals. Arrows indicate the shortened neurites. (F) The length of ALM-N and PLM-PN in various strains indicated. Double asterisks indicate statistical significance ($p < 0.01$) for the difference between *rpm-1* and *mec-15; rpm-1* mutants.
